# Identification and catalogue of viral transcriptional regulators in human diseases

**DOI:** 10.1101/2024.10.06.616669

**Authors:** Citu Citu, Le Chang, Astrid M. Manuel, Nitesh Enduru, Zhongming Zhao

## Abstract

Viral genomes encode viral transcriptional regulators (vTRs) that manipulate host gene expression to facilitate replication and evade immune detection. Nevertheless, their role in non-cancerous diseases remains largely underexplored. Here, we unveiled 268 new candidate vTRs from 14 viral families. We mapped vTRs’ genome-wide binding profiles and identified their potential human targets, which were enriched in immune-mediated pathways, neurodegenerative disorders, and cancers. Through vTR DNA-binding preference analysis, 283 virus-specific and human-like motifs were identified. Prioritized Epstein-Barr virus (EBV) vTR target genes were associated with multiple sclerosis (MS), rheumatoid arthritis, and systemic lupus erythematosus. The partitioned heritability study among 19 diseases indicated significant enrichment of these diseases in EBV vTR-binding sites, implicating EBV vTRs’ roles in immune-mediated disorders. Finally, drug repurposing analysis pinpointed candidate drugs for MS, asthma, and Alzheimer’s disease. This study enhances our understanding of vTRs in diverse human diseases and identifies potential therapeutic targets for future investigation.

## Introduction

Viruses pose a global threat to human health by inducing various cellular effects that often result in infectious and non-infectious diseases, including cancer and immune-mediated disorders. More than 200 viral species, including DNA and RNA viruses, infect humans (Knipe 2013). After infection, virus may integrate into the human genome. So far, over 77,000 virus integration sites (VISs) have been curated (Tang et al. 2020). Viruses primarily contribute to human diseases by encoding viral transcriptional regulators (vTRs). These vTRs interact with nucleic acids either directly (in this case, termed as viral transcription factors) or indirectly through cofactors to modulate host gene expression. Their effects are asserted by influencing the transcriptional machinery or chromatin states, altering immune responses, apoptosis, and cell cycle dynamics (Latchman 1993; Liu et al. 2020). These vTRs play a pivotal role in orchestrating the complex interplay between viral and host cellular processes, ultimately shaping the course of infection and disease progression. As vTRs are crucial for establishing viral genome replication and host gene expression, they are promising therapeutic targets due to their distinct sequential and structural differences from host cell proteins (Nečasová et al. 2022).

Numerous assays have identified vTRs in various viral species, including DNA-binding assays, chromatin immunoprecipitation, and perturbation studies (Lu et al. 2012; Anastasiadou et al. 2019; Liu et al. 2020). Recently, a vTR census was generated by a meta-analysis of viral proteins from various viruses, with evidence of nucleic acid binding, genome targets, secondary structure, and biological function (Liu et al. 2020). Using BLASTp search algorithm, the authors identified 419 vTRs across diverse viral proteomes of DNA and RNA viruses categorized into 20 families. Then, a high-throughput approach validated the vTR’s effector domains from this census and recognized their effect on gene expression when recruited to reporter genes (Ludwig et al. 2023). This study identified the transcriptional regulatory domains of 117 viral proteins, comprising 87 activation domains and 106 repressor domains. The paired yeast one-hybrid assay identified 113 protein pairs between human transcription factors (TFs) and vTRs, arising from protein-protein interactions between them (Berenson et al. 2023). These interactions have been reported to exhibit both cooperative and antagonistic roles, underscoring the noteworthy influence of vTRs on the regulatory elements of the human genome. Despite these notable advances, the comprehensive identification of vTRs across a broad spectrum of viral species using experimentally verified virus regulators and advanced bioinformatic methodologies has not been done yet.

vTRs can directly or indirectly affect the promoters, enhancers, and chromatin regions of human genes (Gordon et al. 2020). For example, several vTRs involving EBNA1, EBNA2, and Zta from Epstein-Barr virus (EBV), LANA and RTA from Kaposi’s sarcoma-associated herpesvirus (KSHV), and E1A from adenovirus can influence gene expression and immune-mediated pathways through genetic and epigenetic changes (Ramasubramanyan et al. 2015; Chiang and Liu 2018). EBNA1, a sequence-specific binding protein, shares structural similarity with DNA-binding domain of LANA, that play important role in maintaining EBV episomes during cell division (Kim et al. 2020). In contrast, EBNA2, EBNA-3s, and EBNA-LP function as coactivators or corepressors by cooperatively binding to human proteins including RBP-jK and EBF1 (Lu et al. 2016). RTA from KSHV is a transcriptional activator that specifies the promoters of target genes by binding directly to or interacting with cellular proteins, thereby acting as a viral E3 ubiquitin ligase which degrades host proteins that block viral lytic replication (Combs et al. 2022). A previous study underscored the functional roles of potential vTR targets in the human genome and identified their co-regulatory functional regions (Liu et al. 2020). Although they are crucial regulatory elements, their full-range genome-wide targets, associated pathways, and links to various diseases are not well understood. Additionally, a comprehensive understanding of the potential DNA-binding preferences of these vTRs, based on genome-wide binding profiles, is yet to be established.

EBV is one of the most common DNA viruses that infect humans. It is mainly involved in Burkitt’s lymphoma, Hodgkin’s lymphoma, and other types of cancer, including nasopharyngeal carcinoma (Kim et al. 2020; Patel et al. 2022). Investigators have been interested in the roles of EBV in distinct autoimmune and neurodegenerative disorders including systemic lupus erythematosus (SLE), Sjögren’s syndrome (SjS), multiple sclerosis (MS), and Alzheimer’s disease (AD) (Chen 2024; Mohammadzamani et al. 2024; Xie et al. 2024). The regulation of EBV vTRs on host target genes in immune-mediated and neurodegenerative diseases during EBV infection is critical but not much clear. The roles of vTRs within the human genome may be modulated by the host genetic makeup, including natural genetic variations (Liu et al. 2020; Rüeger et al. 2021). EBNA2 has been shown to selectively bind genomic loci associated with multiple autoimmune and neurological diseases, influencing disease etiology through allele-dependent mechanisms (Harley et al. 2018; Viel et al. 2024). The association of single nucleotide polymorphisms (SNPs) with different EBV vTRs binding sites and their effects on downstream target genes need to be further investigated.

To bridge these gaps, this study aimed to delineate the roles of vTRs in the human genome and their impacts on diverse diseases through modulating host genes’ expression. We focused on identifying vTRs by applying an ensemble method using experimentally verified vTRs from diverse viral proteomes. Genome-wide chromatin immunoprecipitation sequencing (ChIP-seq) datasets were analyzed to construct vTR-specific binding profiles in the human genome to identify target genes located near these peaks for each vTR. The DNA-binding preferences of these vTRs were identified using a deep learning-based method to underline the virus-specific motifs in the human genome. ChIP-seq profiles were integrated with RNA-seq expression data from EBV virus-infected cells to prioritize the targets of EBV vTRs. Stratified linkage disequilibrium (LD) score regression analysis was performed to quantify the contribution of vTR-binding site annotations to the heritability of complex traits across the genome. Drug repurposing analysis of EBV targets was conducted in a condition-specific manner, aiding the identification of potential drug targets relevant to viral infection. This study provides a holistic understanding of the complex interplay between vTRs and host biology, thereby advancing our knowledge of the contributions of viruses to various human diseases.

## Results

### Ensemble approach detected over 350 vTRs in human viruses

The integrative workflow for identifying vTRs and their effects on the human genome is summarized in Figure 1. The vTRs were identified using an ensemble approach (Chen et al. 2016). that integrated five homology detection tools, each weighted by their accuracy. A previous study identified vTRs using BLASTp, thus has limitations in identifying remote homologs, specifically, with less than <30% identity (Kilinc et al. 2023). The current approach utilized five distinct tools to identify vTRs based on their homology with experimentally validated 212 vTRs, aimed to capture both high- and low-similarity homologs (Table S1). Each tool was assigned a weight based on its accuracy in identifying homologs in a benchmark dataset. The derived weights were multiplied by the min-max normalized score for each homolog and the aggregated scores were retrieved. Notably, the aggregated score performed better than the other five methods in identifying true homologs in a benchmark dataset (Figure 2A). Subsequently, Youden J-statistics calculated 0.076 as the optimal threshold for separating true and false homologs in the benchmark dataset (Figure 2B). This optimal threshold was applied to the vTR homologs, and redundant protein sequences were clustered and manually examined based on their families. A total of 600 vTRs were identified, including 388 homologous vTRs and 212 previously known vTRs (Table S1). Among these 388 identified vTRs, 268 are newly discovered, while 120 overlap with previously reported vTRs (Liu et al. 2020) (Table S1). These 600 vTRs correspond to 191 different viruses (species), including vTRs identified in 97 new viruses (species) such as Bocavirus, Akhmeta virus, and Alston virus

**Figure 1:**
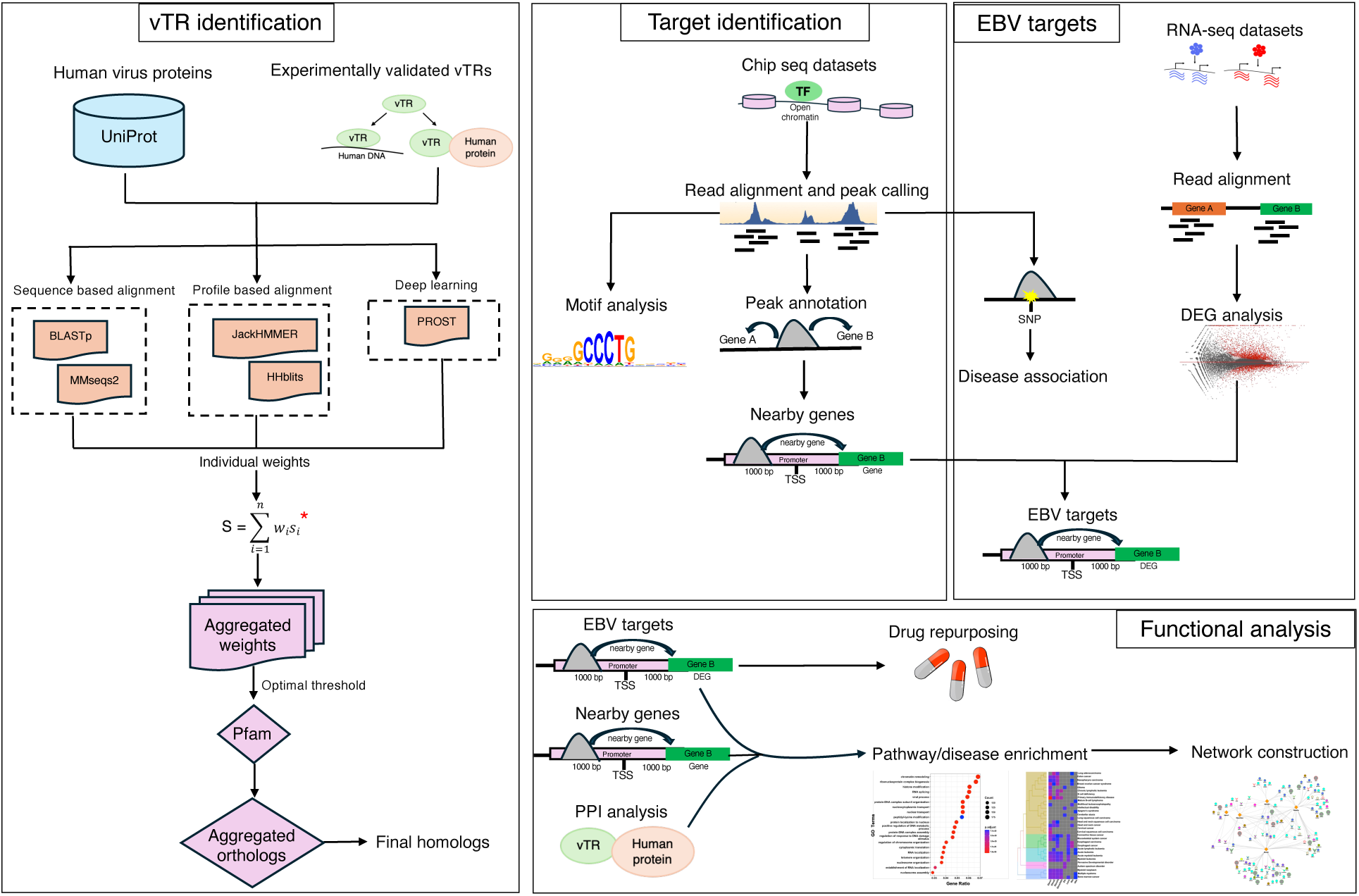
Workflow for viral transcriptional regulator (vTR) identification, characterization, and feature analysis, followed by genetic and drug repurposing analysis. *Chen et al. (Chen et al. 2016).

**Figure 2:**
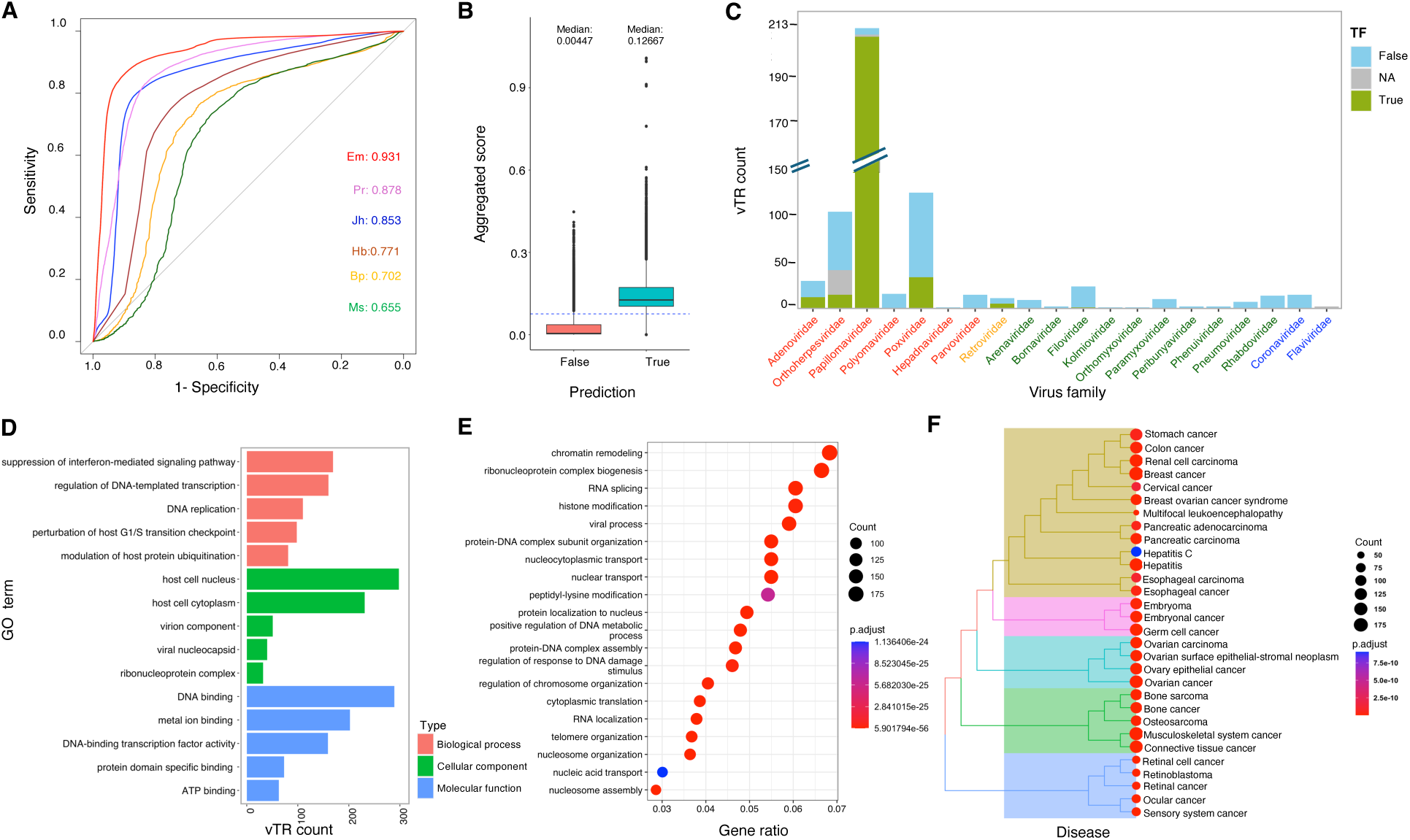
Summary of identification of viral transcriptional regulators (vTRs), their distribution, and functional roles. (A) Receiver operating characteristic (ROC) curve displaying the area under the ROC curve (AUC) values of the six methods in descending order: Em: ensemble method, Pr: PROST, Jh: JackHMMER, Hb: HHblits, Bp: Diamond-BLASTp, and Ms: MMseqs2. (B) Boxplot illustrating the number of true positive and false positive homologs with their aggregated scores in the benchmark dataset. The blue dashed line indicates the cut-off (0.076) used to classify the homologs. (C) Distribution of vTRs across viral families. The color of the labels on the x-axis represents virus types: red (dsDNA), orange (ssRNA-RT), dark green (ssRNA (-)), and blue (ssRNA (+)). TF: transcription factors. NA: not assigned. (D) Top five Gene Ontology (GO) terms for vTRs in each of three GO categories: biological process, cellular component, and molecular function. (E) Gene ontology (GO) enrichment of vTR interactors in the human genome. (F) Disease enrichment of vTR interactors. The clustering of the disease terms was based on pairwise similarities. (Table S1). The vTRs of human viruses are classified among 20 families and the types of genetic material in these families can be classified as ssDNA (+), dsDNA, ssDNA, ssRNA (−), and ssRNA-RT (Figure 2C; Table S1). Their lengths varied from 71 to 3,011 bp, with the smallest length associated with viruses from the *Polyomaviridae* and *Coronaviridae* families, and the highest length classified to viruses from the *Flaviviridae* and *Orthoherperviridae* families. The *Papillomaviridae* family has the highest number of vTRs (213), followed by *Poxviridae* (122), *Orthoherpesviridae* (105), *Adenoviridae* (30), and *Filoviridae* (24). Notably, EBV had the highest number of vTRs (19), followed by KSHV (18), human cytomegalovirus (14), Monkeypox virus (12), Cowpox virus (12), Variola virus (11), and Ectromelia virus (11). These vTRs can be classified into 98 Pfam families, indicating their diverse roles in the human genome. vTRs can modulate host gene expression by directly binding to nucleic acids or interacting with human proteins/TFs. To evaluate whether vTRs act as TFs or bind to DNA using specific domains, a deep-learning-based strategy was employed. This approach identified a number of regulatory entities in vTRs, with the highest number of TFs linked to the *Papillomaviridae* family. Among the 600 identified vTRs of human viruses, 276 (46%) were predicted to act as TFs (Table S1).

The Gene Ontology (GO) terms retrieved from UniProt for these vTRs provided insights into their potential functional roles in the human genome (Figure 2D). The top biological process involves the suppression of interferon-mediated signaling pathways, helping viruses evade the host immune system by escaping immune surveillance and clearance. The vTRs involved in the interferon-mediated signaling pathway of the host primarily correspond to the human papillomavirus (E6 and E7), adenovirus (E1A), and immunodeficiency virus (Tat), which were previously reported to play a role in interferon signaling (Castro-Muñoz et al. 2022). The other top biological processes involve regulation of transcription, DNA replication, modulation of host protein ubiquitination, and perturbation of the G1/S transition checkpoint. The involvement of vTRs in these key cellular processes underscores their intricate role in manipulating the host physiology for viral propagation and pathogenesis (Bagga and Bouchard 2014). Almost half of these vTRs were localized in the host cell nucleus and cytoplasm, probably facilitating their interaction with key cellular machinery. This probably enable them to modulate diverse biological processes that are crucial for viral replication and evasion of host immune responses. Other cellular components, such as virion component, viral nucleocapsid and ribonucleoprotein complex, may help in viral assembly inside the host cell. The primary molecular functions of these vTRs include DNA binding, metal ion binding, DNA-binding transcription factor activity, protein domain binding, and ATP binding activity. These findings further validated their roles in hijacking the host cellular machinery, modulating gene expression, and evading host immune responses. Such activities ultimately promoting viral replication and infection by binding directly to DNA or other human proteins that can regulate gene expression. The identification of new vTRs in various viral species and the resolution of their functions underline their potential impact on gene regulation and immune evasion mechanisms in the host.

### Human interactors of vTRs identified by protein-protein interaction analysis

To elucidate vTR’s effect at the protein level, we performed protein–protein interaction (PPI) analysis to identify their affected human proteins in response to virus infection. In total, we found 6,079 interaction pairs between 90 vTRs and 3,172 human proteins, which involved 250 human TFs (Table S2). Most of these interacting vTRs belong to the *Orthoherpesviridae* (31), *Papillomaviridae* (21), *Retroviridae* (5), *Polyomaviridae* (5), *Coronaviridae* (5), and *Adenoviridae* (5) virus families. These vTRs largely interact with human TFs categorized in the TBP, TP53, and ZNF families, that are crucial for transcription, cell cycle checkpoints, apoptosis, and RNA metabolism during viral infections (Hajikhezri et al. 2020). GO enrichment analysis of human interactors of vTRs indicated their involvement in chromatin remodeling, nucleosome organization, nucleocytoplasmic transport, translation, DNA replication, and RNA splicing. The result supported that vTRs are involved in diverse biological processes and pathways in the genome, epigenome, transcriptome, and proteome (Figure 2E; Table S2). Other human interactors include proteins involved in multiple immune-mediated pathways, transport, protein folding, and degradation. Interestingly, vTRs were found to interact with distinct immune-mediated proteins, including CXCL8, S100s, HLAs, NOTCH, CXCR4, and interleukins. These results highlight their important roles in immune-mediated processes (Scott N. Mueller 2008). We next examined the pathways affected by these interactions for DNA and RNA virus’s vTRs separately. As the result, DNA virus vTRs largely impacted translation-based processes, such as peptide chain elongation and translation initiation, whereas RNA virus vTRs primarily affected mRNA transport and transcription-based processes (Figure S1).

Disease enrichment analysis of these interactors suggested their involvement in diverse types of cancer, including stomach cancer, nasopharynx carcinoma, breast cancer, bone cancer, and retinoblastoma (Figure 2F). In addition, autoimmune diseases, including SjS, psoriasis, and SLE, were also enriched (Table S2). PPI analysis underscored key proteins and pathways influenced during viral infection, shedding light on the potential of these vTRs to contribute to diverse cancers and autoimmune diseases.

### Potential targets of vTRs elucidated using genome-wide binding analysis

To emphasize the genomic effects of vTRs, whether through direct binding to gene promoters or interaction with human TFs, we utilized available ChIP-seq datasets. This allowed us to analyze vTRs’ target genes, affected pathways, and DNA-binding preferences in the human genome. After quality control of 58 available ChIP-seq datasets of vTRs, we obtained 27 high-quality datasets for further downstream analysis (Table S3). The identification of peaks in high-quality datasets using the ChIP-AP pipeline provided a wide range of vTR peaks, which was not possible to capture using a single-peak caller. Subsequently, the peaks from the same vTR were aggregated based on consensus peak calling, thereby identified genome-wide binding of these vTRs in the human genome. In total, we identified the peaks from eight vTRs, two from KSHV (LANA and RTA), one from human cytomegalovirus (HBZ), five from EBV (EBNA1, EBNA2, EBNA3a, EBNA3c and Zta), and a complex of three vTRs from EBV (EBNA3abc). The total number of peaks identified for these vTRs were as follows: 35,120 for LANA, 7,583 for RTA, 5,580 for HBZ, 23,096 for EBNA1, 46,500 for EBNA2, 7,779 for EBNA3a, 36,031 for EBNA3c, 13,140 for EBNA3abc, and 54,084 for Zta. Notably, approximately 80% of the peaks of these vTRs were present near the transcription start site (TSS) of the genes, more specifically, near the -500 bp to +500 bp region for all vTRs except Zta. This evidence supported their regulatory activity in the promoter of human genes (Figure 3A). Peak annotation revealed that their presence was predominantly enriched in the intronic regions in addition to the promoters (Figure 3B). This implies that these vTRs may also act as distal enhancers of gene expression and harbor SNPs that are mainly located in intronic regions (Liu et al. 2020; Nair et al. 2021). The shared number of peaks between these vTRs may indicate their consensus regulation in the human genome. Notably, the limited number of shared regulatory regions between these vTRs probably indicate their distinct regulation in the genome (Figure 3C). The Jaccard similarity between their regulatory regions indicated that there was still some correlation between the regulatory regions of EBNA2, EBNA3s (EBNA3a, EBNA3c, and EBNA3abc), and RTA. These vTRs are transcriptional activators that exert their effects by indirectly binding to the human genome (Wang et al. 2015). The overlapping regions targeted by at least half of these vTRs within the gene promoters identified 236 regulatory regions distributed across the genome (Figure S2A; Table S3). Remarkably, the promoters of *TBC1D1* and *PIK3C2A* genes were regulated by at least seven of these vTRs, indicating that these genes possibly be the key regulatory hubs that are influenced by multiple viral factors (Figures S2B and S2C). Furthermore, these regions exhibited distinct cis-regulatory signals as annotated by The Encyclopedia of DNA Elements (ENCODE), further supporting the potential regulatory mechanisms in the region overlapped by the vTRs. Target genes for the vTRs were identified based on the presence of their peaks near the transcription start site (±1 kb), emphasizing the direct influence of the presence of vTRs on gene expression (Table S4). The highest number of target genes were identified for EBNA2, LANA, and EBNA3c, demonstrating their broad regulatory roles in numerous biological processes (Figure 3D, Table S4). Almost 90% of their target genes are protein-coding, with some being long non-coding RNAs and non-coding genes, underscoring their impact on a broad range of genes (Figure 3D).

**Figure 3:**
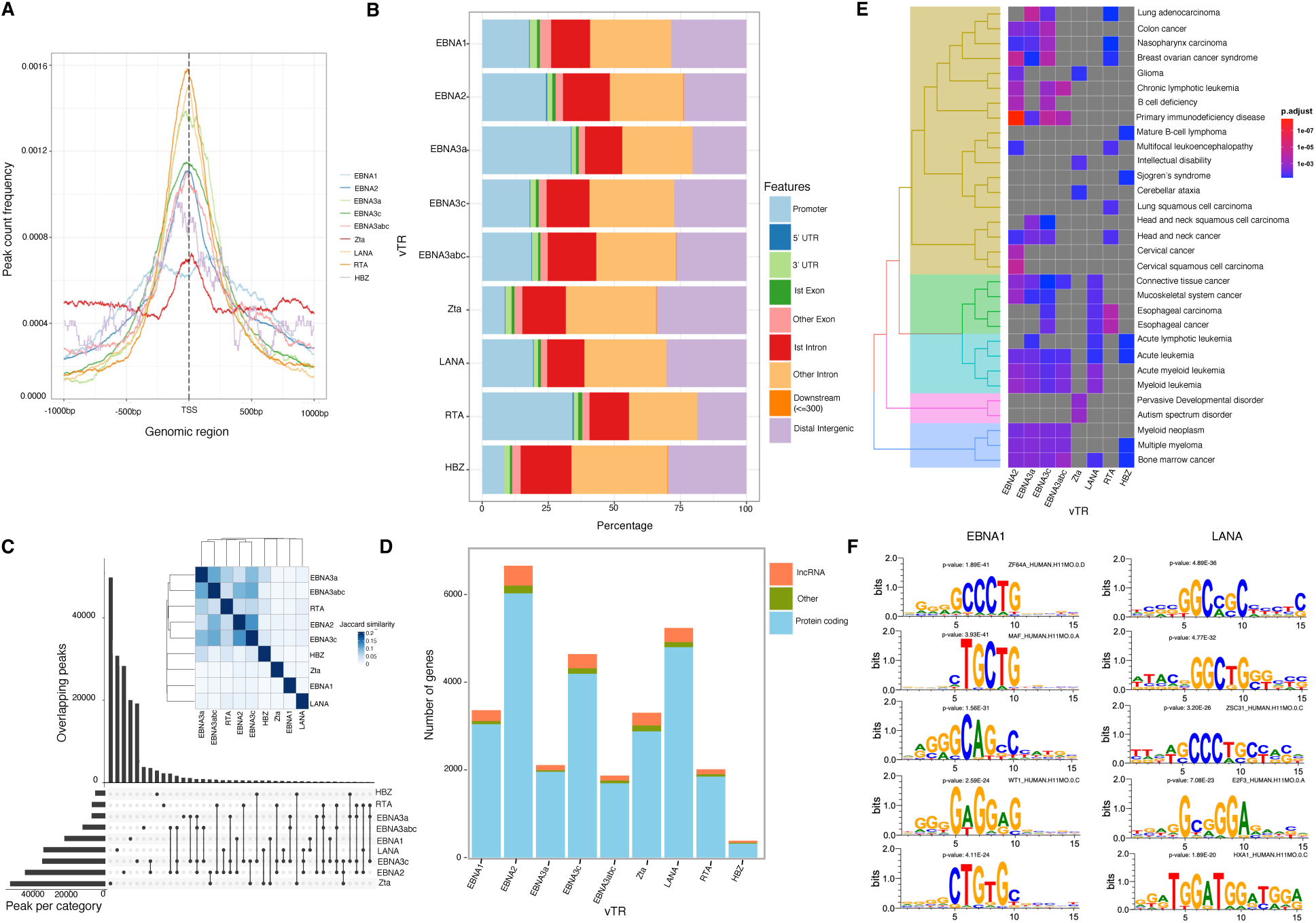
vTR-specific genomic loci, their distribution, associated target genes, and enriched motifs. (A) Distribution of peaks for vTRs near the transcription start site (TSS). (B) Bar stack representing the feature distribution of the vTR peaks in ten genomic regions. (C) UpSet plot showing overlap of peaks between pairs of nine vTRs. The heatmap on the top right corner shows Jaccard similarity between their regulatory regions. (D) Number of genes with TSS located in the +1/-1 kb region near the vTR peaks and their functional roles. (E) Disease enrichment of potential target genes regulated by each vTR. The clustering of the disease terms was based on pairwise similarities. (F) Top five motifs enriched in EBNA1 and LANA peaks with their p-values. Similar motifs in humans were identified using the HOCOMOCO v12 and HOCOMOCO v11 databases with q-values <0.05.

As expected, disease enrichment analysis of their target genes indicated their involvement in diverse types of cancer, consistent with the known oncogenic potential of viral proteins (Figure 3E). This includes lung adenocarcinoma, B-cell acute myeloid leukemia, and nasopharyngeal carcinoma. Common types of cancer among the target genes of vTRs include connective tissue cancer, multiple myeloma, myeloid leukemia, acute leukemia, bone marrow cancer, and stomach cancer, pinpointing the influence of vTRs on a broad range of cancers. Notably, Hodgkin lymphoma (targeted by EBNA2 and EBNA3a) and non-Hodgkin lymphoma (targeted by EBNA2, EBNA3a, and EBNA3abc) genes were regulated by EBV vTRs. This signifies the potential of EBNA2 and EBNA3s in mediating the most common type of cancers associated with EBV infection (Table S4). Additionally, the genes associated with hepatitis B were regulated by EBNA2 and EBNA3a, implying their role in liver injury. Furthermore, the target genes were implicated in distinct immune-mediated and brain-related diseases, including autism spectrum disorder (targeted by Zta), brain ischemia (targeted by EBNA2, EBNA3a, and EBNA3c), SLE (targeted by EBNA2 and EBNA3c), primary immunodeficiency disease (targeted by EBNA2, EBNA3a, EBNA3c, and EBNA3abc), and SjS (targeted by HBZ). These results revealed the broader impact of vTRs on immune system regulation and neurological health in addition to cancer (Table S4).

GO enrichment analysis of vTR target genes underscored their involvement in GTPase binding, GTPase regulator activity, actin binding (targeted by EBNA2, EBNA1, EBNA3abc, Zta), cadherin binding, transcription factor binding, nuclear receptor binding (targeted by EBNA2, EBNA3c, Zta, RTA), phospholipid binding (targeted by EBNA2 and Zta), and ubiquitin-like protein ligase binding (targeted by EBNA2, EBNA3c, LANA, RTA) (Figure S2D; Table S4). These findings highlight the diverse molecular functions of vTRs that affect key cellular processes at the genomic level, including signal transduction, cytoskeletal organization, and gene regulation. The most enriched terms included ATP hydrolysis activity, GTPase regulator activity, nucleoside-triphosphatase regulator activity, transcription corepressor activity, DNA-binding transcription factor binding, cadherin binding, and transcription coactivator activity, emphasizing the leading role of these vTRs in modulating cellular metabolism, structural integrity, and transcriptional networks (Figure S2D). These findings indicated various roles of vTRs, such as their effect on cell-cell adhesion processes that facilitate viral invasion into host cells, transcription factor-related processes to promote virus replication and transcription, and energy metabolism to aid viral spread and hijack cellular mechanisms.

### Genome-wide DNA binding preferences of vTRs identified virus-specific motifs

The vTRs can have different binding preferences compared to human TFs in the genome; therefore, we applied a deep learning-based method to assess the DNA binding specificities of vTRs. This analysis is crucial for understanding their regulatory mechanism in the human genome (Haque et al. 2022). The motifs of these vTRs were preferentially identified by taking the 101 bp flanking region from the peak summit. There were 74 motifs identified for EBNA1, 66 for EBNA2, two for EBNA3a, one for EBNA3c, five for EBNA3abc, 92 for Zta, 37 for LANA, 6 for RTA, and none for HBZ (Figure 3F; Table S5). To determine the corresponding motifs for human TFs, these vTR motifs were scanned against the human TF motif databases with a q-value < 0.05 (Table S6). In addition to identifying motifs corresponding to known human TF motifs, we also identified motifs that did not match any motifs known for human TFs. Our findings indicated that vTRs may have the potential to bind to genomic sites distinct from those targeted by human TFs. For all these vTRs, we identified previously uncharacterized motifs that did not correspond to any known human TF binding sites (Table S5). Notably, the EBNA1 motifs were recognized by VEZF1 (11), ZFX (11), KLF3 (12), SP4 (14), SP2 (15), and SP3 (15), emphasizing that EBNA1 recognizes the motifs of these human TFs and potentially functions with them (Table S6) (Pei et al. 2020). Moreover, 12 EBNA2 motifs were recognized by COE1 (EBF1), that has been previously advised to carry out key metabolic processes together with EBNA2 (Beer et al. 2022). The EBNA3s motifs were recognized by nine IRFs, including IRF4, IRF2, IRF6, and IRF8, found to physically interact with these through B cell development during apoptosis (Banerjee et al. 2013). Zta was recognized by JUN (3) and FOS (4) transcription factors, including FOSB, FOS, JUND, and JUNB that can activate various signaling pathways (Zhang et al. 2016). LANA from KSHV binds with ETV4 and ETV5, thereby reflecting LANA’s role in affecting hematopoiesis. RTA motifs have been found to bind to diverse zinc finger proteins, including ZNF425, ZNF281, and ZNF436. These results suggested that RTA plays a role in modulating gene expression at various levels. Interestingly, the majority of these vTR motifs were recognized by VEZF1 (25), SP3 (24), SP2 (23), JUNB (23) and FLI1 (22) TF motifs, that are known for their roles in diverse types of cancer (Table S6). Our results detailed the distinct DNA-binding preferences of vTRs compared to human TFs, in addition to co-association. Overall, these findings revealed putative regulatory mechanisms and potential therapeutic target sequences in viral-associated diseases.

### EBV vTR targets prioritized through genomic and transcriptomic data integration

The association of EBV vTRs with immune-mediated and neurodegenerative diseases was identified and is shown in Figure 4A. To prioritize the targets of EBV vTRs, we integrated the ChIP-seq and RNA-seq datasets from EBV. In total, we identified 3,092 differentially expressed genes (DEGs), including 1,662 upregulated and 1,430 downregulated, for EBV in virus-infected versus control cells (Table S7). The regulatory potential of EBV vTRs on human genes was identified using ChIP-seq peaks and RNA expression, revealing 1,816 potential targets (Table S8). The EBNA1 targets included 267 upregulated and 192 downregulated genes; EBNA2 targets included 726 upregulated and 531 downregulated genes; EBNA3c targets included 466 upregulated and 341 downregulated genes; EBNA3abc targets included 182 downregulated genes; and EBNA3a targets included 247 upregulated and 189 downregulated genes, and no target was identified for Zta (Figures 4B and 4C, Table S8). The Jaccard similarity between shared genes of EBV vTRs revealed an overlap of EBNA2, EBNA3a, and EBNA3c targets, implying that they may be involved in the same type of mechanism in causing human diseases (Figure 4D). GO enrichment of EBV vTR target genes suggested their involvement in varied activities, including cyclin-dependent serine/threonine kinase (EBNA2, EBNA3a, EBNA3c), helicase activity (EBNA2), RNA binding (EBNA2), serine/threonine inhibitor activity (EBNA3a), and transcriptional repressor activity (EBNA3c, EBNA3abc) (Figure 4E). The serine/threonine activities of EBNA2 and EBNA3s pinpoint their role in cell proliferation, differentiation, and apoptosis. No significant enrichment was identified for EBNA1 targets, indicating that it evidently regulates different sets of genes when compared to other EBNAs (Figure 4E, Table S8).

**Figure 4:**
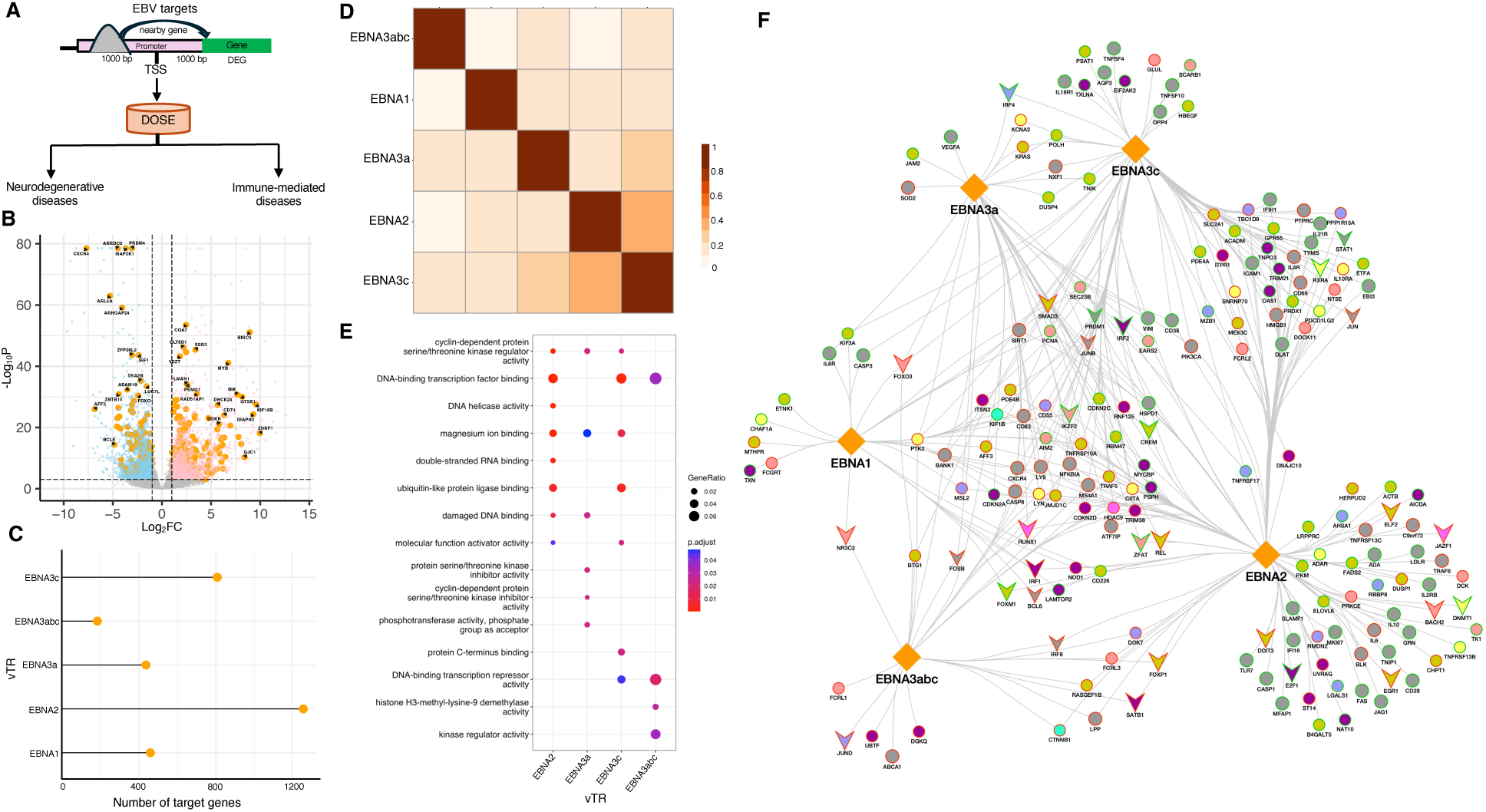
Epstein-Barr virus (EBV) vTR targets and their roles in autoimmune diseases. (A) Workflow illustrates the prioritization of EBV targets and their enrichment in immune-mediated and neurodegenerative diseases. (B) Volcano plot displays the prioritized EBNA1 targets with absolute log2FC >1 and q-value cut-off <0.001. FC: fold change. Upregulated genes are denoted in pink, downregulated genes in blue, and EBNA1 prioritized targets in orange. (C) Number of prioritized targets identified for each EBV vTR. (D) Jaccard plot shows the target genes shared by EBV vTRs, providing insights into overlapping regulatory mechanisms across five EBV proteins. (E) GO enrichment analysis of EBV vTR targets compared with their varied functions. (F) Network representation of EBV vTR targets and their involvement in diverse autoimmune diseases. The node color is below. Cyan: multiple sclerosis (MS); yellow: systemic lupus erythematosus (SLE); peach: Hashimoto’s disease; dark yellow: psoriatic arthritis; blue: myasthenia gravis; pink: rheumatoid arthritis (RA); purple: Sjögren’s syndrome (SjS); and grey: multiple autoimmune diseases. V shape nodes: transcription factors. Orange diamonds: EBV vTRs. Node border color: red denotes downregulated genes and green denotes upregulated genes.

We performed disease enrichment analysis, underlining the potential roles of EBV vTRs in the regulation of a variety of diseases (Figure 4A). The enrichment of EBV vTR targets revealed their significant enrichment in different diseases, with the top associations including non-Hodgkin lymphoma, Hodgkin lymphoma, Burkitt’s lymphoma, and T-cell lymphoma (Table S8). Additionally, these enrichments include neurodegenerative diseases such as Huntington’s disease (HD) and AD, autoimmune diseases involving myasthenia gravis, SLE, rheumatoid arthritis (RA), psoriasis, MS, and Hashimoto’s disease, as well as other immune-mediated disorders such as vitiligo, asthma, primary sclerosing cholangitis (PSC), and allergic rhinitis (AR) (Table S8). As previous studies have predominantly explored the relationship between EBV infection and cancer, we specifically investigated the associations between EBV vTRs, and immune-mediated and neurodegenerative diseases here (Figure S3A). Network generation of autoimmune disease genes and EBV vTRs revealed 183 genes with 352 connections between them (Figure 4F). Specifically, the center of the network represents the genes targeted by multiple vTRs, including major downregulated autoimmunity genes comprising *CIITA, CXCR4*, *LYN*, and *LY9*. These genes function in tolerance of self-antigen and regulation of immune state, and their downregulation can lead to various autoimmune conditions (Caspar et al. 2022). In addition, we emphasized some of the upregulated autoimmune genes, including *VEGFA*, *PRDM1*, *FOXM1*, *CD226*, and *AIM2*, known for their overexpression under different autoimmune conditions (Guo et al. 2022). Notably, we identified 30 TFs in this network, including PRDM1, FOXM1, FOS, FOXP1, and IRF8. These TFs are known to play important roles in autoimmune reactions (Zhou et al. 2022). This implies that EBV vTRs play a significant role in downregulating immune-tolerance genes and activating hyperactive genes, potentially leading to manifold autoimmune diseases. In addition to identifying the role of EBV vTRs in autoimmune diseases, we also elucidated their contribution to neurodegenerative and other immune-mediated disorders (Figures S3B and S3C). We identified genes such as *SIRT1*, *ENO1*, and *IDE* as EBV vTR targets that are important for AD, as well as interleukins, including *IL4* and *IL6*, crucial for immune-mediated diseases (Tian et al. 2023).

Furthermore, we investigated the potential functional roles of genes targeted by EBV vTRs in these diseases. GO enrichment analysis revealed that the top GO terms in autoimmune diseases, except MS, were related to the regulation of cytokine production, leukocyte activation/proliferation, and leukocyte cell-cell adhesion, implying that leukocyte adhesion may be the hallmark of autoimmune responses during EBV infection (Figure S4) (Fan and Sun 2022). MS genes were enriched in interleukin production, neuronal death, and lymphocyte proliferation, reflecting the characteristic MS-related inflammatory and neurodegenerative processes regulated by EBV vTRs. Moreover, the terms in other immune-mediated diseases were largely enriched in T-cell proliferation and activation, crucial for diseases such as asthma and vitiligo (Larché et al. 2003; Sherif S Awad 2021). Target genes related to neurodegenerative diseases, including AD and HD, were found to be enriched in neuronal death and response to oxidative stress, which are key causes of these disorders (Chen et al. 2012). Broadly, the prioritized targets for EBV were found to be involved in numerous diseases other than cancer, underscoring the potential of EBV vTRs to cause a wide range of diseases by affecting immune-mediated signaling and pathways.

### vTR binding sites showed significant enrichment in disease heritability

To elucidate the genetic association of a spectrum of immune-mediated and neurodegenerative diseases to EBV vTRs binding sites, we developed an integrated analysis workflow for quantifying the contribution of the vTR binding site annotations to the heritability of these traits (Figure 5A). Using Stratified Linkage Disequilibrium Score Regression (sLDSC), we demonstrated that genomic annotations corresponding to EBV vTR binding sites significantly contribute to the heritability of distinct immune-mediated and neurodegenerative diseases, as illustrated in Figure 5B. This analysis encompassed five individual vTRs and a complex of three vTRs (EBNA1, EBNA2, EBNA3a, EBNA3abc, EBNA3c, and Zta) from EBV, and assessed their impact on 20 diseases. Of note, type 2 diabetes (T2D) was included as a negative control, because its classification is typically outside of immune-mediated or neurodegenerative categories. The heatmap in Figure 5B displays the coefficient z-scores for each vTR-disease pair, with statistical significance marked by asterisks (** q-value < 0.05, * q-value < 0.1; Figure 5B).

**Figure 5:**
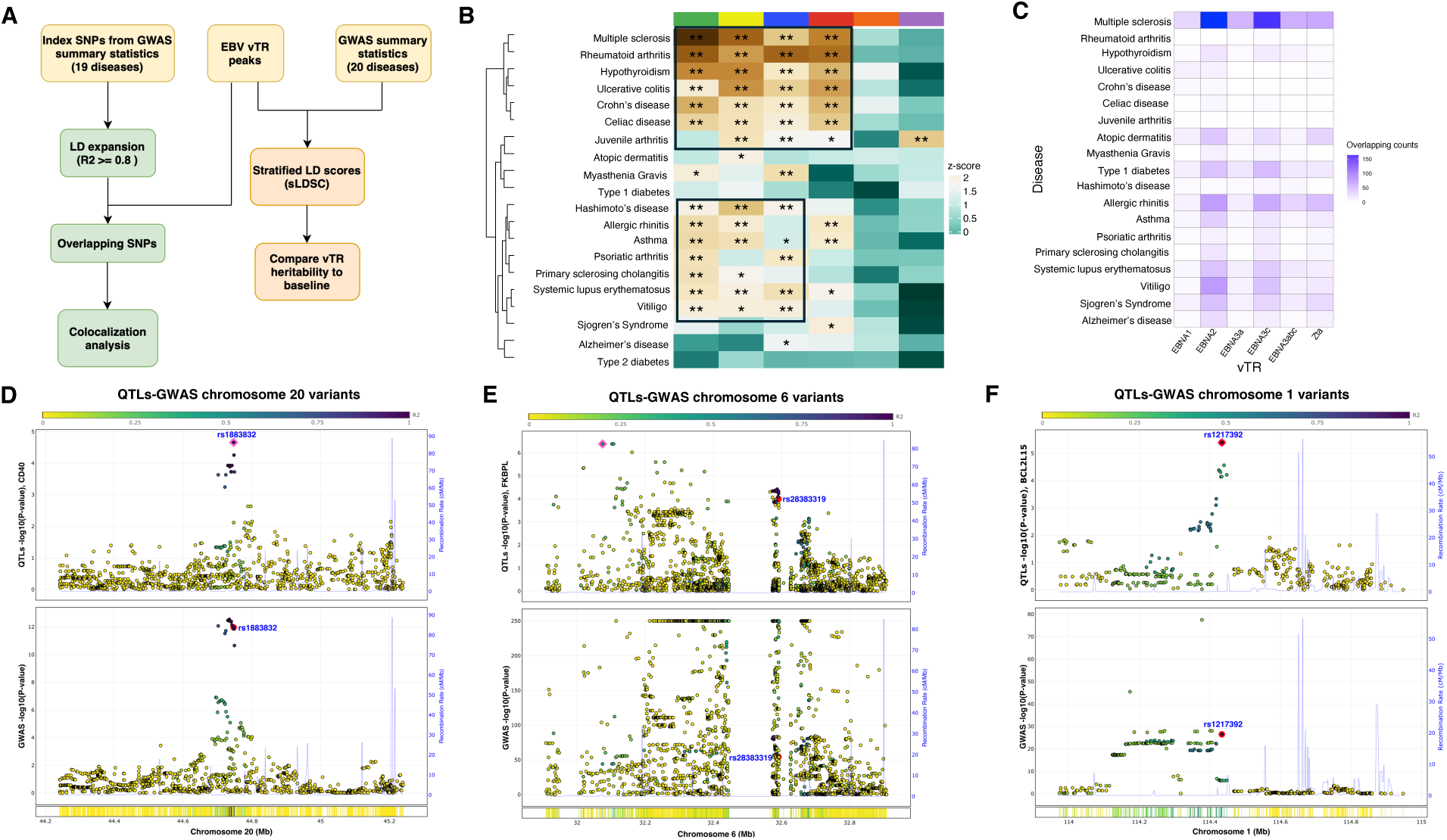
Impact of single nucleotide polymorphisms (SNPs) associated with immune-mediated and neurodegenerative diseases on Epstein-Barr virus (EBV) vTRs’ binding sites. (A) Workflow for identification of the genetic association of EBV vTRs with distinct immune-mediated and neurodegenerative diseases. (B) Enrichment of genome-wide association studies (GWAS) SNPs from immune-mediated and neurodegenerative diseases on EBV vTR peaks as computed by linkage disequilibrium (LD) score regression. Green, EBNA3c; yellow, EBNA3a; blue, EBNA2; red, EBNA3abc; orange, EBNA1; and purple, Zta. (C) Number of SNPs from 19 diseases overlapped with the EBV vTR binding sites. (D) Multiple sclerosis GWAS-expression quantitative trait loci (eQTL) colocalization analysis shows the SNP rs1883832 locus near the *CD40* gene in cortex tissue. (E) Rheumatoid arthritis GWAS-eQTL colocalization analysis shows the rs28383319 locus near the *FKBPL* locus in whole blood tissue. (F) Hypothyroidism GWAS-eQTL colocalization shows the rs1217392 near the *BCL2L15* gene in whole blood tissue.

The EBV vTR binding sites showed widespread enrichment in these diseases, indicating their potential regulatory roles. EBNA2 and EBNA3s binding sites were significantly enriched in multiple autoimmune diseases including MS, RA, Hypothyroidism, SLE, Hashimoto’s disease and psoriatic arthritis, consistent with the previous reports of linking EBV infection to these autoimmune diseases (Harley et al. 2018). Moreover, their binding showed enrichment in other immune-mediated diseases, namely, ulcerative colitis (UC), celiac disease (CD) and Crohn’s disease (CrD), vitiligo, asthma, AR, and PSC, as supported by disease enrichment analysis of their targets. Based on these results, there may exist a cooperative association between EBNA2 and EBNA3s in causing these diseases. EBNA1 binding sites did not show any enrichments in any of these diseases, reflecting its distinct regulation from other EBNAs. The Zta (BZLF1), associated with EBV lytic reactivation, binding sites significantly enriched in juvenile arthritis indicating a potential role for viral reactivation in this disease.

Some diseases showed consistent enrichment across multiple vTR-binding sites. For example, MS had significant enrichment across EBNA2 and EBNA3s vTR binding sites, providing strong genetic support for the long-hypothesized link between EBV and MS pathogenesis (Aloisi et al. 2023). In contrast, diseases such as atopic dermatitis, SjS, and type 1 diabetes (T1D) showed little to no significant enrichment across the vTR-binding sites, suggesting a potentially lower impact of these vTRs on their genetic architecture. Interestingly, AD showed a unique pattern, with moderate enrichment of EBNA2 binding sites. This observation implies a possible, previously underexplored, link between EBV and neurodegenerative processes. This hypothesis warrants further investigation. Our findings align with recent research that revealed a significant genetic overlap between AD and immune-mediated diseases, signifying a shared neuroinflammatory basis that could potentially involve viral factors linking EBV (Enduru et al. 2024). Of note, T2D was included as a negative control in our analysis. Its lack of association with vTR-binding sites supported the specificity of our findings for immune-mediated and certain neurodegenerative diseases.

To further validate these findings, we examined the overlap between disease-associated SNPs and vTR-binding sites (Figure 5C, Table S9). This analysis revealed patterns that largely corroborated with the sLDSC results. MS showed the most striking overlap, with a high number of disease-associated SNPs coinciding with the binding sites of multiple vTRs, particularly EBNA2 and EBNA3c. This finding provides additional support for the robust genetic link between EBV infection and MS pathogenesis. Consistent with the sLDSC analysis, AR, SLE, and vitiligo also showed substantial overlap across multiple vTR-binding sites. The SNPs overlapping within the vTR peaks were analyzed for their linkage with EBV-prioritized targets within ±500 kb regions, specifically from SNPs to the TSS of the associated genes. This analysis revealed connections between the peaks harboring SNPs linked to MS, SLE, and EBV vTR targets (Figure S5A). This convergence of evidence strengthens the hypothesis that common viral regulatory mechanisms underlie numerous immune-mediated, specifically autoimmune, conditions.

### Genome-wide association studies (GWAS) SNPs from three diseases colocalized with expression quantitative trait loci (eQTL)

To further investigate the potential mechanisms occurring in the vTR peaks, we performed co-localization analysis of GWAS SNPs and eQTL signals. Specifically, the top 10 SNPs were selected for this analysis based on their GWAS p-values for MS, RA, and hypothyroidism, respectively, as these diseases demonstrated the highest heritability with EBV vTRs (Figure 5B). For MS, we included five relevant tissues: cerebellum, cortex, spinal cord, whole blood, and EBV-transformed lymphocytes. For RA and hypothyroidism, we included EBV-transformed lymphocytes and whole blood as these are the primary tissues affected in these diseases. This analysis was performed on the ezQTL server, which implements two colocalization methods: eCAVIAR (eQTL and GWAS Causal Variant Identification in Associated Regions) (Hormozdiari et al. 2016) and HyPrColoc (hypothesis prioritization for multi-trait colocalization) (Foley et al. 2021). Significant colocalization was defined as GWAS-eQTL pairs with posterior probabilities greater than the thresholds recommended by the developers of eCAVIAR and HyPrColoc (see Methods). We identified distinct genes with eQTLs associated with SNPs in the vTR-binding regions (Table S9). In MS, eight out of ten top SNPs were associated with eQTLs in at least one of the analyzed tissues. Similarly, we validated eight SNPs for RA and five SNPs for hypothyroidism. Importantly, we identified ten HLA genes colocalized with these disease SNPs, further supporting the genetic associations and potential immunological mechanisms involved in vTRs binding. In MS, we identified GWAS-eQTL colocalization within the region of the lead SNP rs6074022 (chr20:44740196), affecting the expression of *CD40* in both the cortex and whole blood, as shown in Figure 5D and Figure S5D, respectively. Interestingly, *CD40* receptors are critical for the initiation and sustainment of the inflammatory response, a process linked to viral mechanisms in MS (Manuel et al. 2023). Furthermore, this region binds to EBNA3abc (Figure 5D). In RA, we pinpointed GWAS-eQTL colocalization within the lead SNP rs9268645 (chr6:32408527) region, affecting *FKBPL* in whole blood (Figure 5E). The *FKBPL* regulates inflammation and vascular integrity. Additionally, this region is a binding site for EBNA1, possibly regulating *FKBPL* (Figure 5E). We also identified GWAS-eQTL colocalization in hypothyroidism within the lead SNP rs3811019 (chr1:114471583) region, affecting the expression of *BCL2L15* in the whole blood (Figure 5F). The expression of Bcl-2 family genes determines the fate of cells undergoing apoptosis and was implicated in hypothyroidism (Singh et al. 2003). The colocalization of rs3811019 (chr1:114471583; binding site of EBNA2) with *BCL2L15* in whole blood may signify its crucial role in hypothyroidism (Figure 5F). Similarly, we underscored MS SNP-associated regions that colocalized in disease-related tissues with essential genes implicated in MS (Figure S5 B-F). These findings further accentuated the importance of EBV vTRs binding in the genetic architecture of these diseases and their potential regulatory roles.

### Potential candidate repurposable drugs targeting EBV vTR-specific genes

Drug repurposing analysis was performed to identify EBV infection-specific drug targets for MS, asthma and AD. Using this, several drug targets that overlapped and found to be enriched with both vTR targets and genetic risk factors were determined (Table S10). Specifically, our analysis identified 33 drug targets for MS, 33 for asthma, and nine for AD, out of a total of 1,816 vTR targets identified during EBV infection. These drug targets pertained to the categories of ‘successful targets’ (approved by the FDA (Food and Drug Administration)), ‘clinical trial targets’, ‘patented recorded’ or ‘literature-reported targets’. The candidate drugs in ‘successful targets’ category along with their targets and associated conditions are shown in Figure 6. The Etrasimod drug was previously identified as a candidate for the treatment of MS by targeting *S1PR1* (Figure 6A). Our analysis suggested additional drug candidates in addition to those previously identified in MS (Sandborn et al. 2023). Elotuzumab has been proposed as a potential drug for MS in a previous study based on Mendelian randomization (Lin et al. 2023). Of note, we identified a potential repurposable drug candidate targeting *ICAM-1*, previously found to be upregulated in patients with MS and associated with EBV infection. A potential drug targeting *CXCR4* has also been identified. *CXCR4* helps in maintaining the latency of EBV in human cells and acts as a stimulatory factor in MS (Wang et al. 2020). The *CD58* locus was associated with MS, and its associated drug, Alefacept, has been implicated in psoriasis, an autoimmune disease that highlights its potential to treat MS (De Jager et al. 2009). The cytokines *IL4R* and *IL6* were previously found to be linked to asthma and as an EBV vTR target; therefore, drugs targeting these genes may be repurposable drug candidates for this disease (Figure 6B). Dupilumab has been shown to reduce severe asthma by targeting *IL-4R* (Castro et al. 2018). Notably, elevated *DPP4* levels in patients with asthma and their potential association with EBV, along with the identified drugs, emphasize a promising treatment option for asthma. Moreover, *SLC23A2* has been implicated in *APOE*-associated cognitive decline, and drugs targeting this gene may be candidates for reducing the risk of AD during EBV infection (Figure 6C). The drug Ursodeoxycholic acid is associated with *SLC23A2*. This drug was found to improve mitochondrial function in patients with AD, underlining its potential to act as a

**Figure 6:**
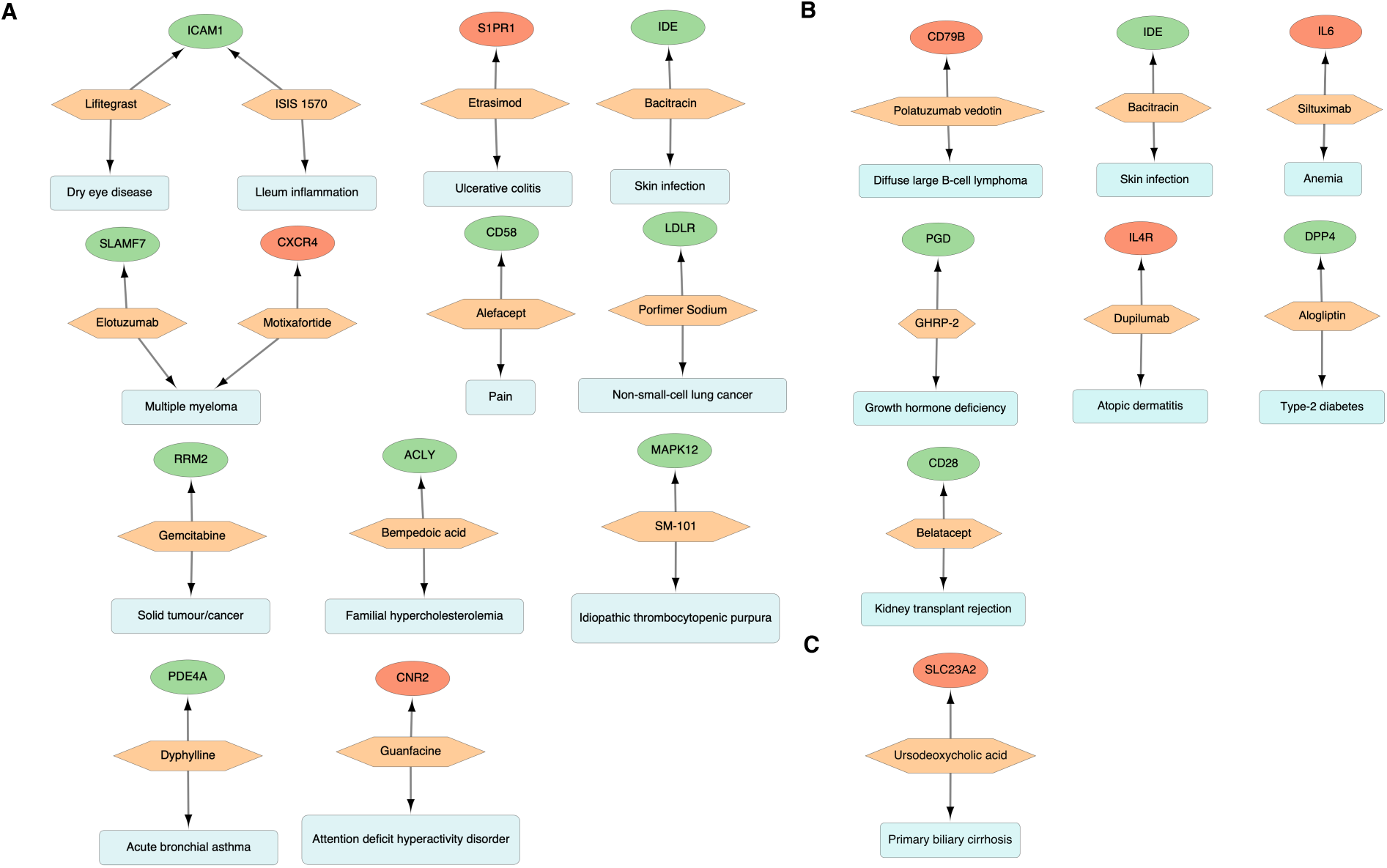
Drug repurposing analysis of Epstein-Barr virus (EBV) vTR target genes. (A-C) Predicted drugs and associated conditions for EBV vTR targets in (A) multiple sclerosis (MS), (B) asthma, and (C) Alzheimer’s disease (AD). Ellipse: target genes. Hexagon: drugs; Rectangle: associated conditions. Red nodes denote downregulated genes, and green nodes denote upregulated genes. candidate drug in AD during EBV infection (Bell et al. 2018). In summary, our analysis pinpointed potential drugs associated with EBV targets for reducing or htreating associated conditions.

## Discussion

In this study, we applied an advanced ensemble approach based on homology detection to identify vTRs across human viral species. We identified a total of 600 vTRs from 191 viruses, including 388 homologous vTRs and 212 previously known vTRs. Moreover, among the 388 identified vTRs, 268 vTRs across 14 viral families were not previously reported. This result not only validated the efficacy of our computational pipeline but also provided promising vTR candidates for future investigation. We next performed comprehensive analyses to catalog vTRs across virus families, including regulators from understudied viruses such as Akhmeta, Alston, and Bocavirus. Furthermore, we studied how these vTRs are associated with leading complex diseases including various cancers, brain disorders, and immune-mediated diseases, and informed potential drug target candidates. Our study significantly expands our understanding of virus-host interactions and their implications for human health.

Our results revealed potential roles of vTRs in modulating key cellular pathways, particularly those involved in immune evasion and cellular reprogramming. By influencing interferon-mediated responses, chromatin modeling, and nucleosome positioning, vTRs exhibit a remarkable capacity to reshape host cellular architecture. Based on our genome-wide binding profiles, vTRs not only preferentially associate with promoter regions but also function as distal enhancers, indicating their extensive regulatory capacity and their interactions with the SNPs linking to the specific disease (Borsari et al. 2021). The major difference between their binding sites might imply their distinct regulatory roles in the genome. Of note, EBNA2 and EBNA3s shared several regulatory sites, reflecting their possibly co-regulatory roles. The identification of target genes based on their peak proximity underscored their pivotal role in regulating protein-coding genes as well as their significant involvement in several types of cancer and immune-mediated diseases, including autoimmune disorders. Interestingly, the genes targeted by EBNA2 and EBNA3s have been implicated for bone marrow cancer, lung adenocarcinoma, nasopharyngeal carcinoma, breast ovarian cancer, multiple myeloma, SLE, SjS, and primary immunodeficiency disease. In summary, the proximity of vTRs regulatory sites near promoters, the diversity of their regulatory roles, and their ability to target disease-related genes highlight the potential of vTRs to regulate a broad spectrum of biological functions under diverse conditions.

We examined the DNA-binding preference of these viral regulators in the human genome using a deep-learning approach. Comparison of vTRs motifs with human TF motifs pinpointed 135 out of 283 virus-specific motifs that did not match any known human motifs. Further investigation of these motifs is warranted. Approximately 53% of the identified vTR motifs are recognized by distinct sets of human TFs, emphasizing their diverse regulatory roles in the human genome. The 62 vTR motifs were found to bind to the VEZF1, SP2, SP3, JUNB, FLI1 and SP1 motifs. This indicated the potential of these DNA sequences as the viral targets in cancer (Li et al. 2015). Deep learning approaches have also been applied to predict virus integration sites and unveil the unique motifs at those sites (Xu et al. 2021; Xu et al. 2023). Understanding the binding preferences of regulators is critical, and the current identification of specific vTR binding sites may pave the way for further clinical research focusing on targeting these sites in various diseases during viral infection.

The integrated genomic and transcriptomic approach prioritized EBV vTR target genes and revealed their potential roles in serine/threonine kinase activity, helicase activity, and DNA binding. This supported the notion on virus-host interactions in cell division and proliferation, but our results provided more specific details. The disease enrichment of EBV vTR targets also linked them to diverse cancer, immune-mediated, and neurodegenerative disease genes. GO enrichment analysis of the target genes in these diseases emphasized that leukocyte adhesion and cytokine formation are key processes regulated by these vTRs in immune-mediated diseases and neuronal death in neurodegenerative diseases. Specifically, EBV vTR targets encompass crucial genes linked to autoimmune diseases, such as *CXCR4*, *ENO1,* and *CIITA* (Devaiah and Singer 2013; Caspar et al. 2022). Overall, the integration of expression with their DNA-binding profiles and prioritization of EBV vTR targets revealed their significant enrichment in immune-mediated pathways and diseases, elucidating key genes implicated in these conditions in EBV infection.

Our genetic analysis of vTR-associated variants in broad diseases provides valuable insight into the complex interplay between viral factors and human health. This is important to find human genetic variants that are more susceptible to the risk of viral infection (Dai et al. 2021). The observed enrichment patterns not only supported existing hypotheses regarding virus-disease associations but also led to innovative perspectives on how vTRs may influence disease susceptibility through their interaction with the host genome. We found the consistent enrichment of EBV-related vTR binding sites across multiple autoimmune diseases, particularly in MS, reinforcing the long-standing hypothesis of EBV involvement in MS pathogenesis (Aloisi et al. 2023). The unexpected association between certain EBV vTRs and diverse diseases, including RA, SLE, UC, CrD, CD, hypothyroidism, and AD, opens new avenues for investigating the role of vTRs in these conditions. These findings underline the intricate relationship between viral elements and human diseases, emphasizing the need for further mechanistic studies to elucidate the specific molecular pathways that mediate these associations. To validate these candidate loci, investigators may perform functional studies, investigate their roles in disease mechanisms, and evaluate their utility as molecular targets for therapeutic intervention or as biomarkers for disease risk prediction. For example, our colocalization analysis of the top SNPs associated with vTR binding in MS, RA, and hypothyroidism revealed 21 eQTLs, including *CD40* for MS. Drug repurposing analysis pinpointed several candidate drugs for MS, asthma, and AD. We identified previously reported drugs such as Etrasimod, as well as the additional candidate drugs that may prevent these diseases during EBV infection.

In summary, large-scale identification of vTRs from 191 viral species provides important resources for future investigation of these regulators in a broad spectrum of human diseases. The identified target genes for vTRs underscored their impact on potential biological processes in human. In addition to confirming previously reported diseases, our study identified different associations between vTRs and complex diseases. The prioritized targets of EBV vTRs discern their involvement in diverse immune-mediated and neurodegenerative diseases such as MS, RA, SLE, and AD, emphasizing the specific roles of EBV vTRs in these conditions. The genetic association also underlined numerous SNPs associated with these diseases and their potential roles in affecting vTR binding and various eQTLs in the human genome. These findings suggest that vTRs play crucial roles in modulating biological processes and disease pathways in diverse viral infections. The exploration of these regulatory proteins and their interactions with the human genome could further provide more insights into viral pathogenesis.

## Methods

### Ensemble method for vTR identification

To systematically identify vTRs from available viral species, a total of 212 experimentally verified viral regulators were collected from previous studies (Liu et al. 2020; Berenson et al. 2023; Ludwig et al. 2023). Next, we collected a dataset of ∼1.5 million reviewed and unreviewed viral protein sequences from the UniProt (https://www.uniprot.org) database by setting human as the host. To identify the robust homologs of these 212 experimentally validated vTRs, five tools, including Diamond BLASTp (Buchfink et al. 2021), MMseqs2 (Steinegger and Söding 2017), JackHMMER (Finn et al. 2011), HHblits (Remmert et al. 2011), and PROST (Kilinc et al. 2023), were employed for the ensemble method, which defined different weights for each tool based on their accuracy. Considering that these tools rely on distinctive principles for homolog identification, using multiple tools can ensure the comprehensive coverage and robust identification of vTRs between virus sequences and the human genome. To define weights for each tool, a benchmark dataset containing 7,329 protein sequences that were classified into 1,824 families and 1,070 superfamilies was retrieved from Astral (https://astral.berkeley.edu) based on the SCOPe database (https://scop.berkeley.edu). The homologs of these 7,329 sequences were assessed within the same dataset using the abovementioned five tools with following parameters, --query-cover 50; --id 50; --evalue 1e^-6^ for Diamond BLASTp, and e-value 1e^-6^ for other four tools. The accuracies of these methods were evaluated using the pROC package (v1.18.4) (Robin et al. 2011) based on true homologs, which were identified when the query and target proteins pertaining to the same family. The weight of each tool was calculated using the following formula:

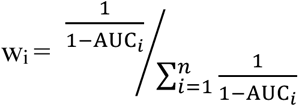

where w_i_ is the weight for tool *i* and AUC_i_ is the area under the receiver operating characteristic curve (AUC) value of each tool.

The calculated weighed and normalized z-scores for each tool were used to calculate the aggregated score, as shown in Chen et al. (Chen et al. 2016).

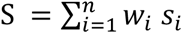

where w_i_ is the weight for tool *i*, s_i_ denotes the normalized score, and S is the aggregated score of identified homologs.

The optimal cutoff for the identification of true homologs based on the aggregated score was calculated using Youden J-statistics (Ruopp et al. 2008), implemented in the pROC package (Robin et al. 2011). The optimal cutoff ascertained by the benchmark dataset was used to elucidate the homologs of 212 vTRs in human viral proteins retrieved from UniProt. The type of genetic material in these viruses was identified using the Virus-Host DB (Mihara et al. 2016).

The InterProScan (Quevillon et al. 2005) tool was used to evaluate the family of the homologs, and their GO terms were retrieved from UniProt. DeepTFactor, a deep learning-based tool, was used to predict TFs from homologous protein sequences (Kim et al. 2021).

### Protein-protein interaction analysis

The experimentally verified interactions between human proteins and vTRs were retrieved from the MINT(Chatr-aryamontri et al. 2007) and HVIDB databases (Yang et al. 2021), containing 48,643 interactions covering 35 virus families and 6,633 virally targeted host complexes aggregated from different databases. GO and disease enrichment analyses of the interacting proteins of vTRs were performed using clusterProfiler (Yu et al. 2012) and DOSE (Yu et al. 2015b) R packages at an q-value cut-off < 0.05. Functional enrichment of interacting proteins of vTRs from DNA and RNA viruses was conducted using gProfiler (Raudvere et al. 2019) at q-value cutoff <0.05. The GO networks were built using the EnrichmentMap (Reimand et al. 2019) pipeline with default parameters in Cytoscape (Shannon et al. 2003) (v3.9.1).

### Analyzing vTR target genes and pathways through ChIP-seq analysis

To identify the genes and pathways targeted by these vTRs in humans, 58 ChIP-seq datasets have been collected from previous studies (Table S2). These datasets were processed using the ChIP-seq Analysis Pipeline (ChIP-AP v5.0) by leveraging multiple peak calling methods (Suryatenggara et al. 2022). This approach recognizes that individual peak callers exhibit distinct specificity and selectivity, leading to a minimal overlap between their results. To ensure the quality of the datasets, normalized strand coefficient (NSC) >1.05 and relative strand correlation (RSC) >0.8 from the phantompeakqualtools (https://www.encodeproject.org/software/phantompeakqualtools/) were set as criteria to remove the low-quality datasets. To retrieve all the associated peaks for each vTR from multiple cell lines, the peaks were merged using bedtools (Quinlan and Hall 2010) *merge* function based on at least 1 bp overlap between the peaks. The macs2 (Zhang et al. 2008) *bdgpeakcall* from the MACS2 tool was used to call the peaks from the merged bed files for each vTR. Blacklist regions were removed from the peaks using the hg38 blacklisted bed file from ENCODE (Abascal et al. 2020). The intersection between the regulatory regions of the vTRs was identified using the *bedtools jaccard* function of bedtools. The nearby genes from the peaks of each vTR were identified using the ChIPseeker package (Yu et al. 2015a), and the peaks residing ±1 kb from the TSS were considered as target genes. To evaluate the pathways and diseases associated with the target genes of the vTRs, clusterProfiler and DOSE packages were used at a q-value cut-off <0.05. The cis-regulatory elements in the candidate promoters were identified using the *SCREEN* web server in ENCODE.

### Motif analysis of vTRs by deep learning

To assess the DNA-binding specificities of vTRs, a deep learning approach based on DNABERT was employed (Ji et al. 2021). The authors demonstrated DNABERT could improve transcription factor binding site (TFBS) analysis when compared to conventional methods with dependence on position-specific weights. Specifically, the pre-trained model of DNABERT was fine-tuned based on the ChIP-seq peaks for each vTR. To this end, we extracted 101 bp (50 bp upstream and 50 bp downstream) sequences around the summit position of the analyzed peaks, thereby labelled as positive sets. The same number of negative sequences (101 bp each) were extracted randomly from the genome without overlapping with any peaks. The total sequences were split with a 70:30 ratio into the training and testing datasets. The fine-tuning of these sequences using DNABERT was executed using the default parameters: learning_rate 2e-4, -- num_train_epochs 5.0, and --weight_decay 0.01. The fine-tuned model for each vTR was used to calculate the attention scores, and the motifs were identified using the *find_motifs.py* script of DNABERT using the parameters --min_len 6, --pval_cutoff 0.001, and --min_n_motif 5. The identified motifs for each vTR were compared against the HOCOMOCO v11 and HOCOMOCO v12 motifs using the TomTom tool in Meme-Suite (Bailey et al. 2015) with the q-value <0.05. The q-value represents the minimum false discovery rate required to include the match.

### RNA-seq analysis and DEG identification from EBV

The EBV vTR targets identified using ChIP-seq peaks were prioritized by the integration of RNA-seq datasets from GSE125974 (Wang et al. 2019) and GSE155345 (Lamontagne et al. 2021), which profiled the expression of human genes in EBV-infected and non-infected B cells. In the first step, FASTQ reads were trimmed to remove low-quality reads and adapter sequences using fastp (Chen et al. 2018), and quality assessment of the trimmed FASTQ reads was conducted using FastQC (https://www.bioinformatics.babraham.ac.uk/projects/fastqc). Subsequently, the trimmed reads were aligned with the human genome reference (GRCh38) using the STAR (Dobin et al. 2013) (v2.7. b). After aligning the reads, a count matrix was generated, representing the number of reads mapped to each gene in the human genome using *featureCounts* in the Subread package (Liao et al. 2014). Gene count matrices were used to call upregulated and downregulated DEGs in virus-infected versus uninfected cells by using DESeq2 (Love et al. 2014) with absolute log2FC >1 and a q-value cut-off < 0.001. Here, FC denotes fold change of gene expression and computes the magnitude of change in gene expression between different conditions.

### Prioritization of EBV vTR targets

The DEGs identified from EBV RNA-seq datasets and the ChIP-seq targets of EBV vTRs were analyzed together to assess the regulatory potential and targets of each vTR using BETA (Wang et al. 2013) at p-value <0.001 in the ±1 kb flanking region of TSS. GO enrichment for the identified EBV vTR targets was performed using clusterProfiler, and disease enrichment was performed using the DOSE R packages with q-value <0.05. A volcano plot was generated using the EnhancedVolcano R package (https://github.com/kevinblighe/EnhancedVolcano) and networks were generated using Cytoscape.

### Assessing the genetic impact of vTRs on disease susceptibility

sLDSC (Finucane et al. 2015) was used to investigate the potential genetic impact of vTRs on susceptibility to immune-mediated and neurodegenerative diseases. This method leverages summary statistics from GWAS and annotated linkage disequilibrium (LD) scores to estimate the disease heritability attributable to specific genomic annotations. We obtained GWAS summary statistics of 20 diseases from GWAS Catalog (Sollis et al. 2023) and GWAS Atlas (Watanabe et al. 2019) databases (Table S11). These included 18 immune-mediated diseases (AR, asthma, atopic dermatitis, CD, CrD, Hashimoto’s disease, hypothyroidism, juvenile arthritis, myasthenia gravis, PSC, psoriatic arthritis, RA, SjS, SLE, MS, T1D, UC, vitiligo), one neurodegenerative disease (AD) and one negative control (T2D) (Table S11). To create vTR-specific annotations, the binding sites of the EBV vTRs, including EBNA1, EBNA2, EBNA3a, EBNAabc, EBNA3c, and Zta, were used to generate custom LD score annotations. These annotations, corresponding to genomic regions potentially regulated by each vTR based on their binding patterns, allowed us to use LDSC to estimate and quantify the heritability of regions associated with each vTR, thereby providing insights into their genetic contributions to disease susceptibility.

### Mapping overlapping SNPs with vTR peaks

Significant SNPs associated with 19 immune-mediated and neurodegenerative traits (excluding the negative control: T2D) were retrieved. To capture the broader genetic context of these SNPs, LD proxy searches were conducted using PLINK (Purcell et al. 2007) (v1.9). Proxy SNPs were defined using an LD threshold of R^2^ ≥0.8, ensuring a comprehensive representation of potentially relevant genetic variants. Here, R^2^ denotes the squared correlation between allelic values at two loci in LD analysis. The binding sites of EBV vTRs were processed to identify the regulatory regions potentially involved in modulating gene expression. These binding sites were formatted as BED files, specifying the genomic coordinates of vTR interactions. The GenomicRanges (Lawrence et al. 2013) package was used to create a genomic representation of both SNPs and vTR peaks, and the *findOverlaps* function was applied to identify SNPs that overlapped with vTR peaks.

### Colocalization of GWAS SNPs and eQTL

We performed colocalization analysis of GWAS and eQTLs signals to validate the genomic loci within the binding sites of EBV vTRs. For this, we identified lead GWAS SNPs within ChIP-seq regions. Considering that the GWAS of MS, RA, and hypothyroidism had strong associations with EBV, we focused on these three diseases here. We performed GWAS-eQTL colocalization analysis using the ezQTL (Zhang et al. 2022) web server and QTL datasets from the Genotype-Tissue Expression (GTEx) Consortium (Lonsdale et al. 2013). This web server employs eCAVIAR (Hormozdiari et al. 2016) and HyPrColoc (Foley et al. 2021) to perform colocalization analysis. The eQTL tissues included in this analysis were chosen by their relevance to the disease. For instance, tissues of the central nervous system were included for MS GWAS-eQTL colocalization, whereas whole blood tissue was included for RA and hypothyroidism analyses. Colocalization was performed with several parameters, including the region of ±500 kb pairs from the lead SNP and the LD reference population of Europeans from the 1000 Genomes project. We used the default parameters in eCAVIAR and HyPrColoc, i.e., posterior probability calculation >0.01 and >0.50, respectively.

### Repurposable drug candidate identification

To investigate potential drug targets and repurposable drug candidates, we first explored vTRs overlapping with genetic risk factors for diseases associated with EBV. To this end, we compiled GWAS for the above-mentioned disease conditions associated with EBV and applied the Multi-marker Analysis of GenoMic Annotation (MAGMA) (de Leeuw et al. 2015) tool to obtain gene-level association scores. MAGMA SNP-wise gene analysis was performed by annotating the SNP-level association scores to the respective gene regions. Subsequently, MAGMA combines the SNP-level p-values using a regression-based framework that considers LD structure among the SNPs. We performed multiple test corrections using the Benjamini-Hochberg (BH) method to obtain the q-values for each gene. To define the genetic risk factors for each disease, we applied a threshold of q-value (BH) < 0.05. After identifying the vTR targets associated with genetic risk factors for each respective disease, we further explored whether these genes overlapped with existing drug target genes annotated in the Therapeutic Target Database (TTD) (Zhou et al. 2024). To ascertain the significance of drug target enrichment, we performed two statistical tests, the Fisher’s exact test and the hypergeometric test. Each test considered the number of drug targets identified for each disease, as well as the number of genes overlapping between the vTR targets and disease-specific genetic risk factors. Specifically, the enrichment tests considered number of genes in the intersection (genes in both the list and TTD database), genes in the list but not in the TTD database, genes in the TTD database but not in the list, and genes in neither the list nor the TTD database. Lastly, we focused on the diseases with significant enrichment of drug target genes, MS, asthma, and AD.

## Supporting information

Number of viral transcriptional regulators (vTRs) in human viruses.

Protein-Protein interactions between vTRs and human proteins, their GO and disease enrichment.

Number of ChIP-seq datasets analyzed with their peak counts and intersecting peaks by at least half of these vTRs in the promoter regions of genes.

Viral transcriptional regulator peaks present in the +/−1 kb region of the transcription start site (TSS) of genes, their GO and disease enrichment.

Number of DNA motifs identified for each vTR with their top motif hits from the human genome.

Number of human TF motifs similar to vTR binding motifs.

Differentially expressed genes (DEGs) for Epstein-Barr virus (EBV).

Prioritized Epstein-Barr virus (EBV) vTR targets based on ChIP-seq and RNA-seq dataset integration and their disease enrichment.

Number of GWAS SNPs overlapping with Epstein-Barr virus (EBV) vTR peaks and their colocalization in MS, RA and hypothyroidism.

Candidate drugs identified for Epstein-Barr virus (EBV) vTRs in multiple sclerosis (MS), asthma, and Alzheimer's disease (AD).

Summary of the Alzheimer's disease and immune-mediated diseases GWAS datasets.

## Acknowledgements

We thank lab members of the Bioinformatics and Systems Medicine Laboratory (BSML) for their valuable discussion. Z.Z. was partially supported by National Institutes of Health grants (R01LM012806, R01CA276513, and U01AG079847). We thank the technical support from the UTHealth Cancer Genomics Core funded by the Cancer Prevention and Research Institute of Texas (CPRIT, RP180734 and RP240610). L.C. was a CPRIT Postdoctoral Fellow in the Biomedical Informatics, Genomics, and Translational Cancer Research Training Program (BIG-TCR) funded by CPRIT (RP210045). A.M.M. was supported by a training fellowship from the Gulf Coast Consortia, on the NLM Training Program in Biomedical Informatics & Data Science (T15LM007093). The funders had no role in the study design, data collection and analysis, decision to publish, or preparation of the manuscript.

## Author contributions

Conceptualization: Z.Z.; methodology: C.C., L.C., A.M.M., N.E., and Z.Z.; formal analysis: C.C., L.C., A.M.M. and N.E.; investigation: Z.Z; resources: Z.Z; data curation: C.C., L.C., A.M.M. and N.E. ; writing – original draft: C.C., L.C. and A.M.M.; writing – review & editing: N.E. and Z.Z.; visualization: C.C., L.C. and A.M.M.; supervision: Z.Z.; project administration: Z.Z.; funding acquisition: Z.Z.

## Declaration of interests

The authors declare no competing interests

## Notes

### Competing Interest Statement

The authors have declared no competing interest.

