## Supplementary material for "Identification and catalogue of viral transcriptional regulators in human diseases": Figure S1-S5

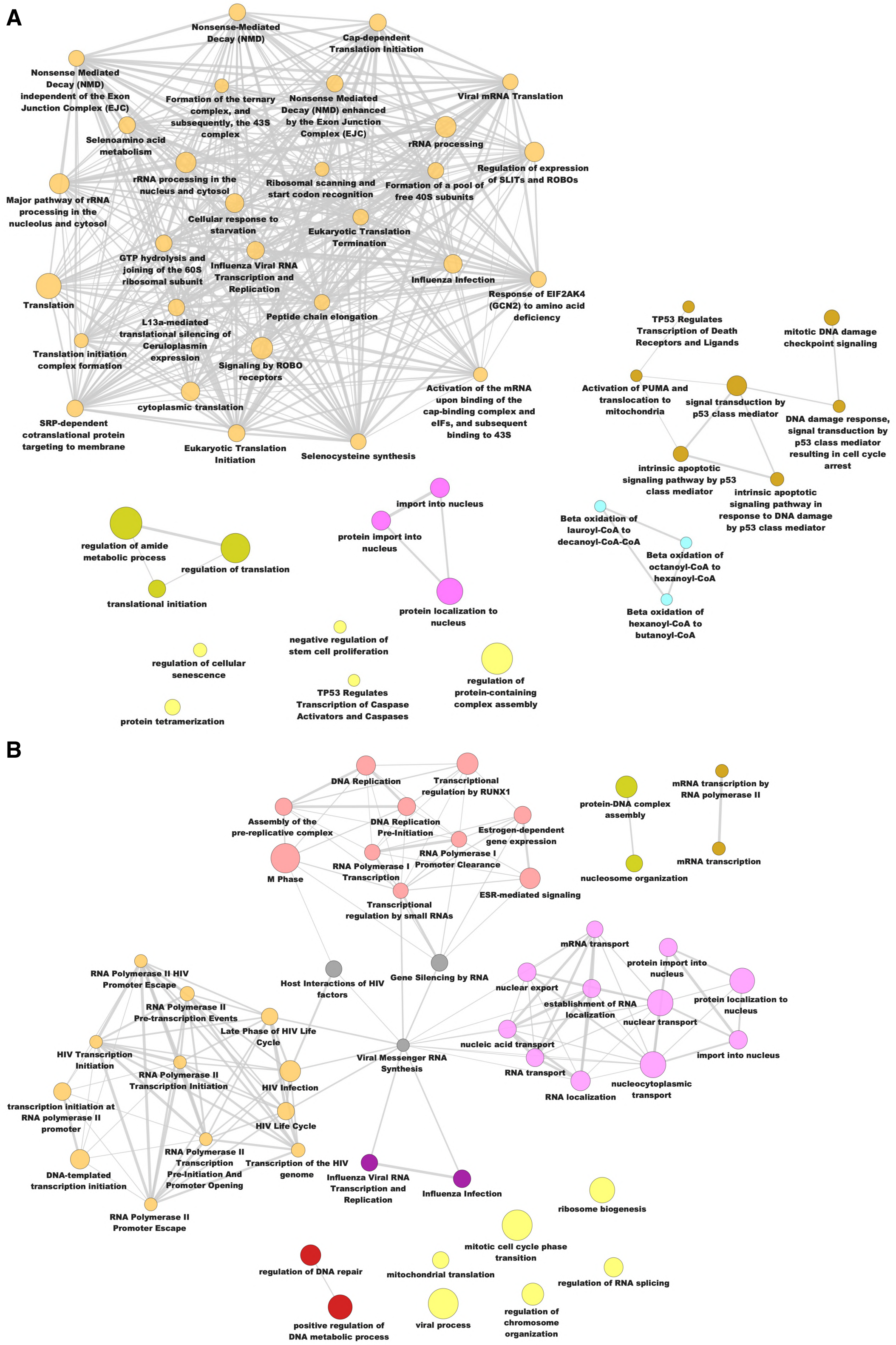


**Figure S1: GO enrichment analysis of vTRs' human interactors.** (A) DNA viruses and (B) RNA viruses.

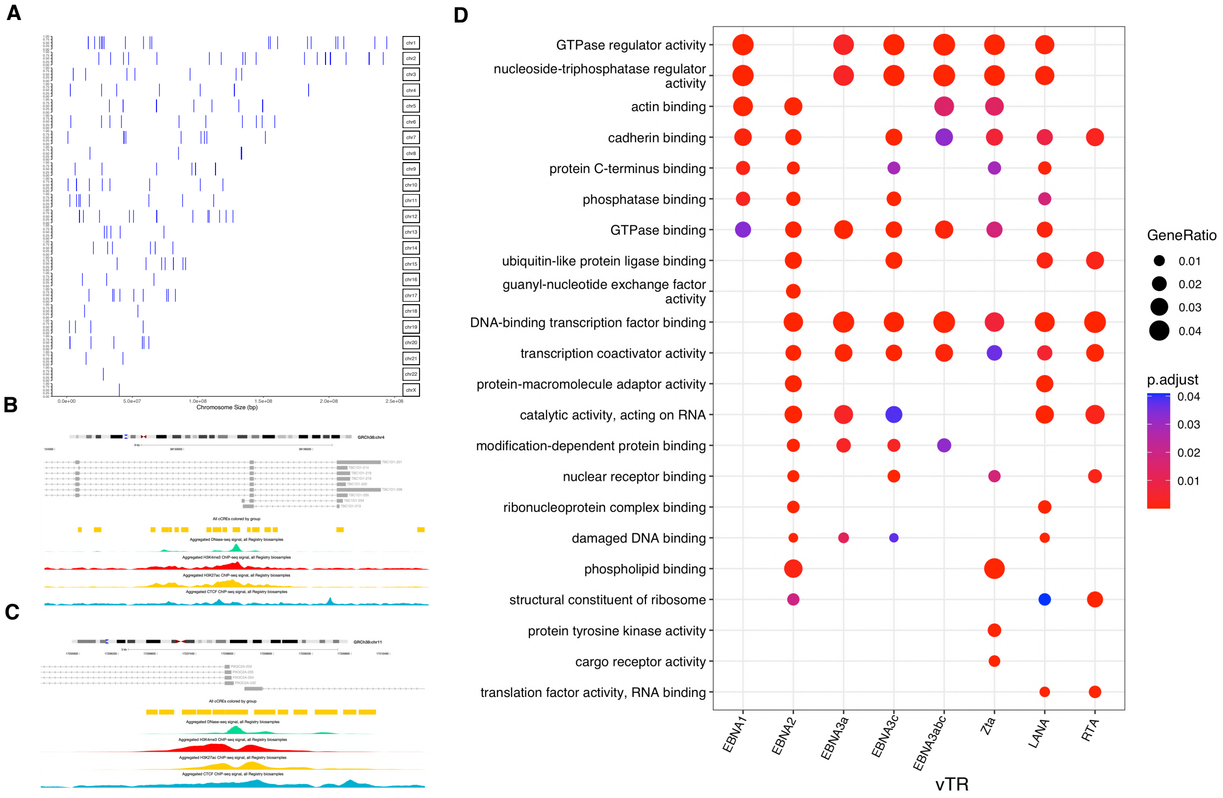


**Figure S2: Genomic distribution of vTR regulatory regions and enriched pathway of the target genes.** (A) Genomic distribution of regions intersected by vTRs showing their numerous distributions on chromosome 1 and 2. (B-C) Association of cis-regulatory elements on the promoter of *TBC1D1* (B) and (C) *PIK3C2A* genes. (D) GO enrichment analysis of vTR targets.

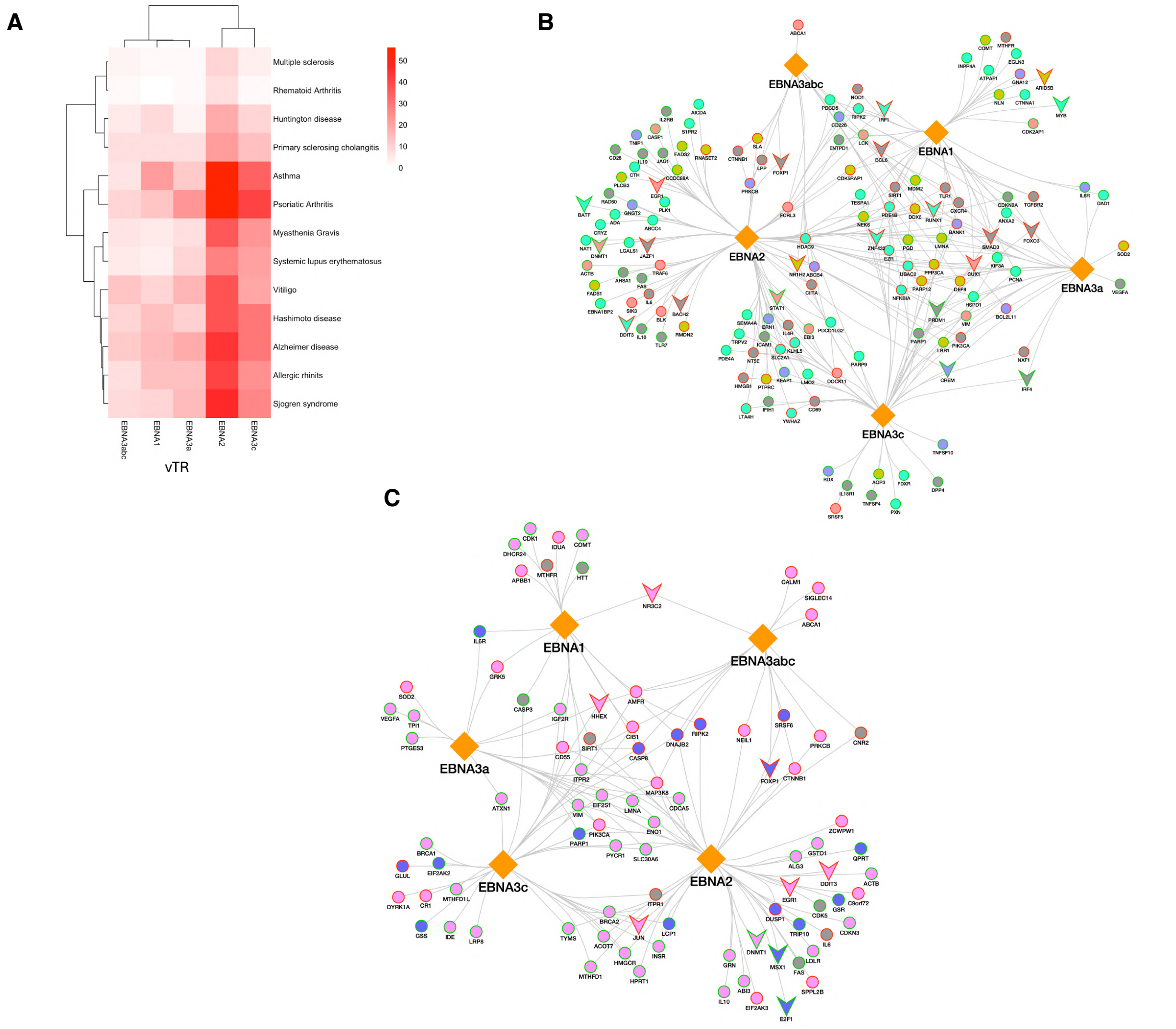

**Figure S3: Epstein-Barr virus (EBV) vTRs' prioritized targets associated with immune-mediated and neurodegenerative diseases.** (A) Number of EBV vTR targets identified in immune-mediated and neurodegenerative diseases. (B) Network representation of EBV vTR targets and their involvement in immune-mediated diseases. Peach: allergic rhinitis (AR); cyan: asthma; blue: primary sclerosing cholangitis (PSC); dark yellow: vitiligo; and grey: multiple immune-mediated diseases. Node border color: red denotes downregulated genes and green denotes upregulated genes. (C) Network representation of EBV vTR targets and their involvement in neurodegenerative diseases. Pink: Alzheimer's disease (AD); blue: Huntington's disease; and grey: both diseases. Node border color: red denotes downregulated genes and green denotes upregulated genes.

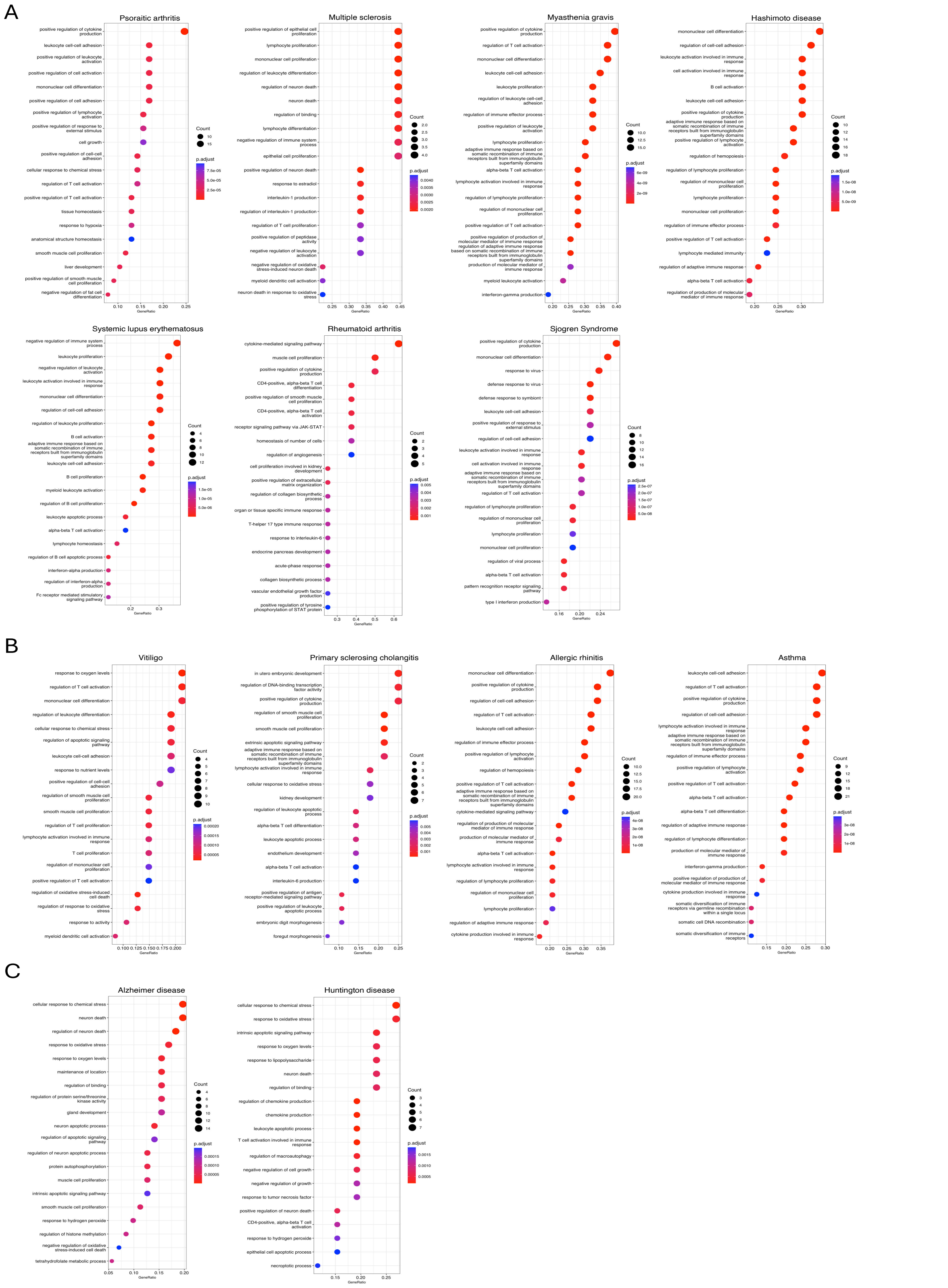


**Figure S4: GO enrichment of Epstein-Barr virus (EBV) vTRs' prioritized targets from different diseases.** (A) Autoimmune, (B) Other immune-mediated, and (C) Neurodegenerative disorders.

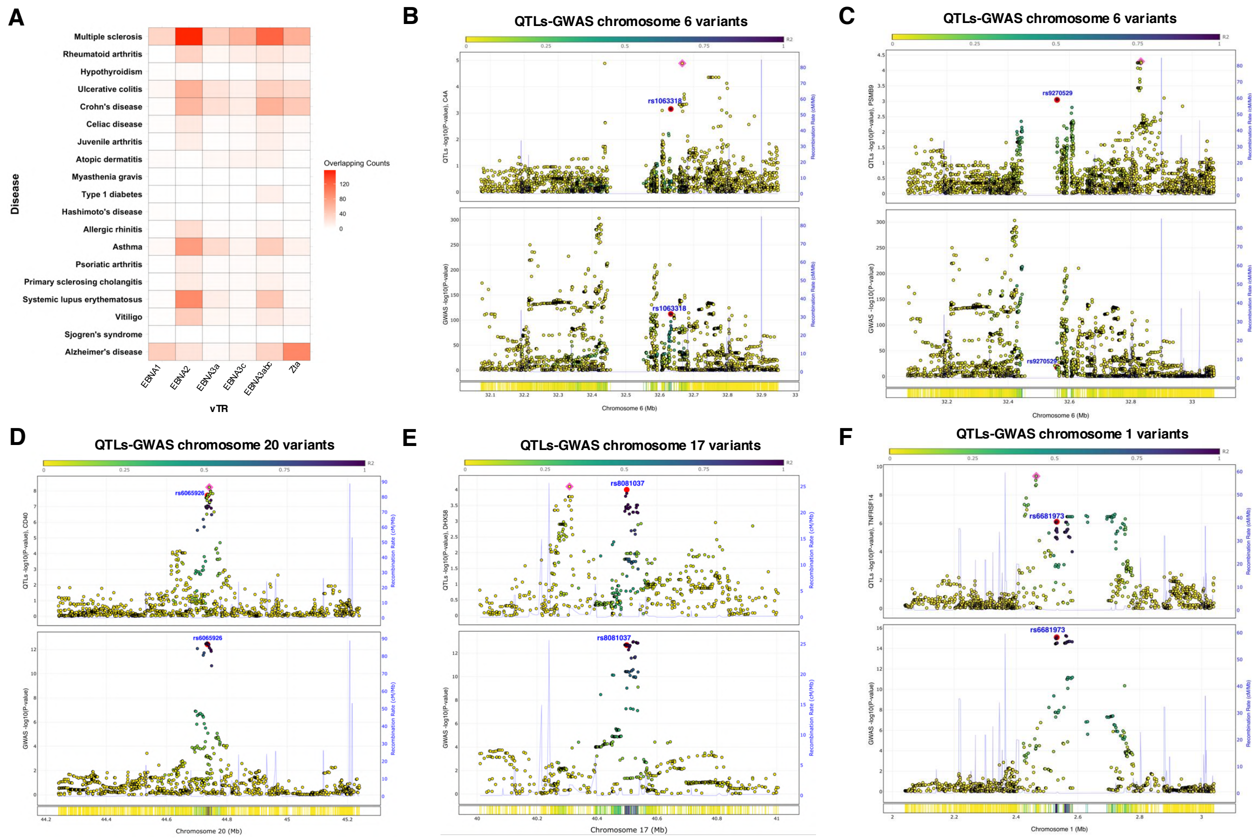


**Figure S5: Impact of disease-related single-nucleotide polymorphisms (SNPs) on target genes.** (A) The number of peaks with disease-related SNPs linked to Epstein-Barr virus (EBV) target genes. (B) Multiple sclerosis (MS) genome wide association studies (GWAS)-expression quantitative trait loci (eQTL) colocalization shows the SNP rs1063318 locus near the *C4A* gene in cortex tissue. (C) MS GWAS-eQTL colocalization shows the rs9270529 locus near the *PSMB9* gene in spinal cord. (D) MS GWAS-eQTL colocalization shows the rs606592 locus near the *CD40* gene in cortex tissue. (E) MS GWAS-eQTL colocalization shows the rs8081037 locus near the *DHX58* gene locus in cortex tissue. (F) MS GWAS-eQTL colocalization shows the rs6681973 locus near the *TNFRSF14* gene in cerebellum.
