## Supplementary material for "Identification and catalogue of viral transcriptional regulators in human diseases": Summary of the Alzheimer's disease and immune-mediated diseases GWAS datasets.

**Table S11**. Summary of the Alzheimer's disease and immune-mediated diseases GWAS datasets, Related to STAR Methods.

| Trait | # SNPs | # cases | # controls | Reference |
|---|---|---|---|---|
| Alzheimer's disease | 12,674,020 | 39918 | 358140 | (Wightman et al. 2021) |
| Allergic rhinitis | 10,321,706 | 22057 | 267250 | (Watanabe et al. 2019) |
| Asthma | 10,599,055 | 44301 | 341521 | (Watanabe et al. 2019) |
| Atopic dermatitis | 11,296,421 | 10788 | 30047 | (Paternoster et al. 2015) |
| Celiac disease | 523,399 | 4533 | 10750 | (Dubois et al. 2010) |
| Crohn's disease | 9,649,131 | 12194 | 28072 | (De Lange et al. 2017) |
| Hypothyroidism | 10,154,468 | 13043 | 231847 | (Watanabe et al. 2019) |
| Primary sclerosing cholangitis | 7,891,603 | 4796 | 19955 | (Ji et al. 2017) |
| Rheumatoid arthritis | 8,747,963 | 14361 | 43923 | (Okada et al. 2014) |
| System lupus erythematosus | 7,915,252 | 7219 | 15991 | (Bentham et al. 2015) |
| Ulcerative colitis | 9,666,299 | 12366 | 33609 | (De Lange et al. 2017) |
| Vitiligo | 8,721,264 | 4680 | 39586 | (Jin et al. 2016) |
| Multiple sclerosis | 8,589,719 | 47,429 | 68,374 | (Patsopoulos et al. 2019) |
| Juvenile arthritis | 7,461,261 | 3,305 | 9,196 | (López-Isac et al. 2021) |
| Myasthenia gravis | 23,768,005 | 1,873 | 36,370 | (Chia et al. 2022) |
| Type 1 diabetes | 25,837,085 | 7666 | 583280 | (Sakaue et al. 2021) |
| Hashimoto thyroiditis | 10,585,639 | 16191 | 552642 | (Sakaue et al. 2021) |
| Psoriatic arthritis | 8,552,612 | 5,065 | 21,286 | (Soomro et al. 2022) |
| Sjögren's syndrome | 25,844,643 | 1599 | 658316 | (Sakaue et al. 2021) |
| Type 2 diabetes | 25,845,070 | 84224 | 583280 | (Sakaue et al. 2021) |
